# No evidence for a special role of language in feature-based categorization

**DOI:** 10.1101/2021.03.18.436075

**Authors:** Yael Benn, Anna A. Ivanova, Oliver Clark, Zachary Mineroff, Chloe Seikus, Jack Santos Silva, Rosemary Varley, Evelina Fedorenko

## Abstract

The relationship between language and human thought is the subject of long-standing debate. One specific claim implicates language in feature-based categorization. According to this view, language resources facilitate object categorization based on a certain feature (e.g., color). Specifically, it is hypothesized that verbal labels help maintain focus on a relevant categorization criterion and reduce interference from irrelevant features. As a result, language impairment is expected to affect categorization of items grouped according to a single feature (low-dimensional categories, e.g., ‘Things that are yellow’), where many irrelevant features need to be inhibited, more than categorization of items that share many features (high-dimensional categories, e.g., ‘Animals’), where few irrelevant features need to be inhibited. In two behavioral studies with individuals with aphasia, we failed to find consistent support for the role of language in low-dimensional categorization. We also collected fMRI data from healthy adults and observed little activity in language-responsive brain regions during both low-dimensional and high-dimensional categorization. Combined, these results demonstrate that the language system is not implicated in object categorization. Our work adds to the growing evidence that, although language may assist in accessing task-relevant information (e.g., instructions), many cognitive tasks in adult brains proceed without recruiting the language system.

## 1. Introduction

The role of language in mediating or augmenting thought is the subject of long-standing debate. According to one view, language is necessary for many cognitive functions, such as math, logic, and thought (e.g., Baldo et al., 2010, 2015; Bermúdez, 2007; Bickerton, 1995; Carruthers, 2002; Darwin, 1871; Dennett, 1994). However, a large body of evidence supports a different view: that language is cognitively and neurally independent from the rest of human cognition. This evidence includes the lack of activity in the language brain regions during non-linguistic tasks that allegedly require language (e.g., Amalric & Dehaene, 2016, 2019; Fedorenko et al., 2011; Ivanova et al., 2021; Monti et al., 2009, 2012), the retained ability of some individuals with aphasia to perform such tasks (e.g., Bek et al., 2013; Benn et al., 2013; Siegal & Varley, 2006; Varley et al., 2005), and variability across cultures in the use of language resources during thought (Kim, 2002). However, the role of language is still contested for one important aspect of human cognition: categorization.

Like other animals, humans can convert rich, multi-dimensional perceptual inputs into a latent lower-dimensional structured representation of the world. Grouping discriminable individual objects and events into classes allows us not only to decide whether some new object/event belongs to a particular category, but also to draw powerful inferences about shared properties from one category member to another (e.g., Mareschal & Quinn, 2001; Mervis & Rosch, 1981; Murphy, 2002; Pearce, 1994; E. E. Smith & Medin, 1981; L. B. Smith & Heise, 1992; Wasserman et al., 1988). In contrast to other animals, humans additionally label individual categories with words—the core building blocks of a powerful communication system that allows us to share complex thoughts with one another. The link between words and learning the structure of new categories has been extensively investigated in infants/children (e.g., Ferguson & Waxman, 2017; Gershkoff-Stowe et al., 1997; Plunkett et al., 2008; Sloutsky & Fisher, 2004; Waxman & Gelman, 2009) and, to some extent, in adults (Brojde et al., 2011; Lupyan et al., 2007; Lupyan & Casasanto, 2015). But how does language affect the process of grouping objects into categories when the category boundaries are already known?

### 1.1 High-dimensional and low-dimensional categories

Before summarizing the key evidence, it is important to introduce a distinction that is theoretically and empirically relevant to this question. Lupyan and colleagues (e.g., Lupyan, 2009; Lupyan & Mirman, 2013; Perry & Lupyan, 2014) distinguish between ‘high-dimensional’ (HD) categories, where members share many features, and ‘low-dimensional’ (LD) categories, where members share one or a few features. HD categories typically correspond to established sets that reflect either the taxonomic (similarity-based) or relational/thematic (co-occurrence-based) structure of the world (Bain, 1864; Mirman et al., 2017). Taxonomic HD categories can often be labeled by superordinate terms such as ANIMALS, FRUIT, or TOOLS. Relational HD categories correspond to common events/scenarios: for example, THINGS YOU TAKE ON A PICNIC or NON-FOOD THINGS FOUND IN THE KITCHEN. For such relational categories, the shared features have to do with typical co-occurrences (e.g., although a fridge and a spatula are quite different, they both co-occur with a large number of kitchen objects, like a stove, pots and pans, a kettle, etc.). In contrast to HD categories, LD categories are more likely to be novel groupings of items that often straddle taxonomic and relational boundaries, such as THINGS MADE OF WOOD or THINGS THAT ARE YELLOW (e.g., things made of wood may include a cupboard, a sledge, and a wooden spoon, and things that are yellow may include a lemon, a yellow hat, and a canary).

Similar distinctions have been made by others, in related literatures. For example, Barsalou (1983) distinguishes between ‘common’ categories, which mirror the correlational structure of the environment, and ‘ad-hoc’, or ‘goal-derived’, categories, which are constructed for a specific goal and are thus often based on a small number of features. Kloos & Sloutsky (2008) and Sloutsky (2010) distinguish between ‘dense’ and ‘sparse’ categories based on the ratio of category-relevant variance to total variance. Members of statistically dense categories share many inter-correlated features that matter for category membership, and members of sparse categories have very few features in common, with many other features varying independently and being irrelevant for category membership. Couchman et al. (2010) contrast family-resemblance categorization, which relies on judgments of overall similarity, considering multiple features in tandem, and criterial-attribute categorization (or ‘rule-based categorization’), which requires adhering to a single-dimensional criterial attribute and suppressing all other, irrelevant dimensions (see also Ashby & O’Brien, 2005). Langland-Hassan et al. (2021) relate the HD/LD distinction to the concrete/abstract distinction, arguing that items in concrete categories have many shared features, whereas identifying items from an abstract category requires generalizing over many irrelevant properties to identify a small set of commonalities.

### 1.2 The LD-specific language recruitment hypothesis

One claim that emerged in the literature in recent years is that language plays a special role in LD categorization (Lupyan, 2009, 2012; Lupyan & Mirman, 2013). The argument goes as follows: categorizing objects into LD/sparse categories is more cognitively costly because features irrelevant to the categorization criterion interfere and have to be inhibited; for instance, when categorizing objects by color, their shape and function have to be ignored. A verbal label (e.g., ‘yellow’) can help maintain focus on the relevant categorization criterion^1^ and reduce interference from irrelevant features. The LD-specific language recruitment hypothesis makes two predictions: i) reduced availability of language resources should lead to a greater disruption of LD compared to HD categorization^2^; and ii) LD categorization should engage the language system to a greater degree than HD categorization.

The first prediction had found some support in the aphasia literature. Some patients with linguistic deficits have been reported to exhibit impairments in non-verbal categorization tasks when the task required focusing on one particular dimension and ignoring other salient dimensions (Cohen et al., 1980; Cohen & Woll, 1981; Davidoff & Roberson, 2004; De Renzi & Spinnler, 1967; Hjelmquist, 1989). Building on these findings, Lupyan (2009) manipulated verbal vs. spatial interference in a dual-task paradigm in neurotypical participants and found that verbal, but not visuo-spatial, interference affected the participants’ ability to decide whether an object belongs to an LD category. In contrast, verbal and visuo-spatial interference had similar (and negligible) effects on HD categorization. Lupyan and colleagues concluded that access to lexical resources (verbal labels) is important for LD categorization.

In a follow-up study, Lupyan and Mirman (2013) directly compared performance on HD and LD categorization in individuals with aphasia and neurotypical controls. Participants were provided with a category label and then had to select from a picture array the subset of objects that belong to the target category (similar to **Figure 1, top**). Performance in the LD condition was lower for both groups, but critically, the HD vs. LD difference was larger in individuals with aphasia, particularly in those with low scores on a picture-naming task.

**Figure 1.**
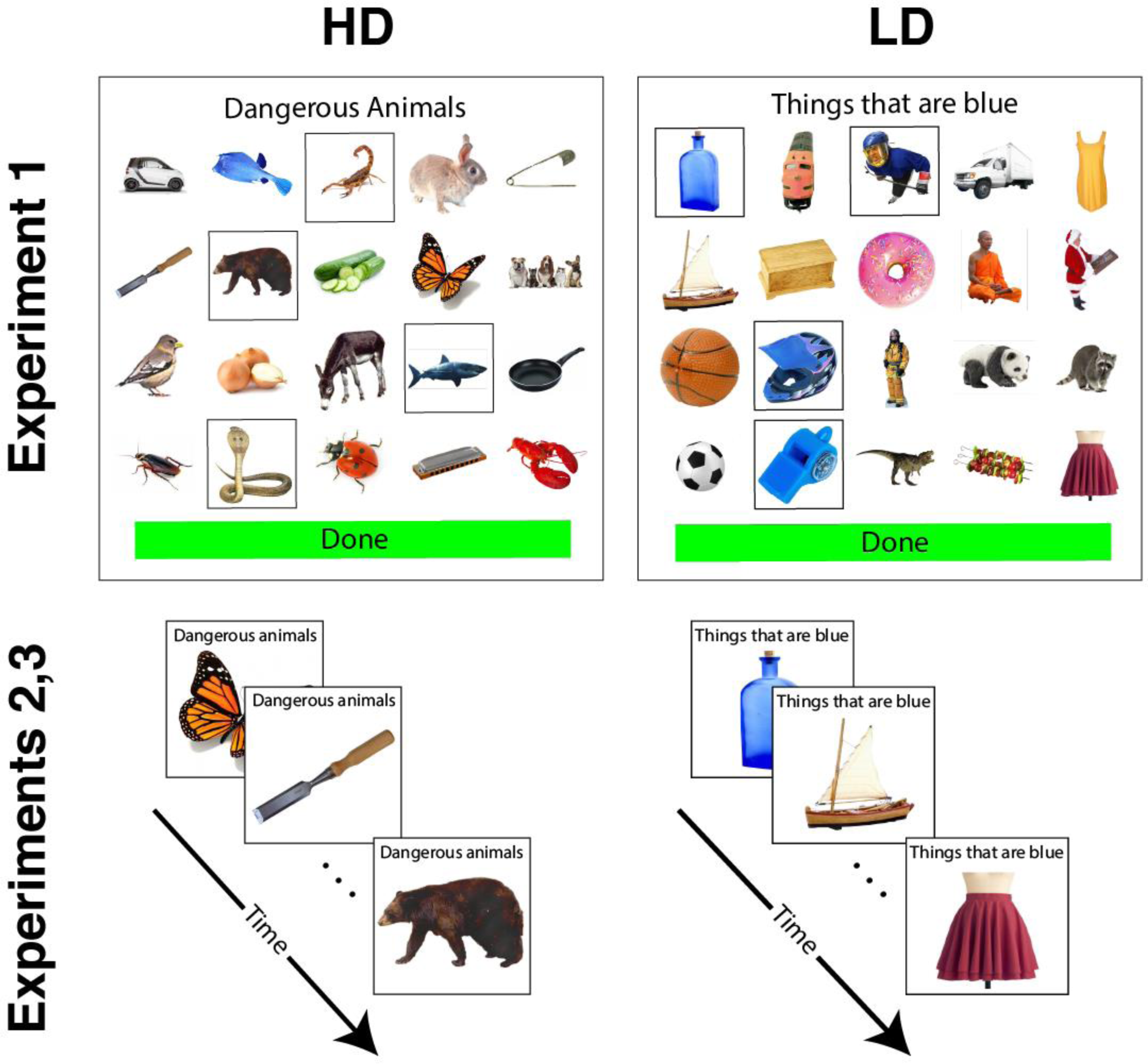
Trial structure in Experiment 1 (top) and Experiments 2 and 3 (bottom). HD – high dimensional category, LD – low dimensional category.

However, evidence from aphasia does not provide uniform support for the LD-specific language recruitment hypothesis. For example, Burger and Muma (1980) found deficits in HD categorization in individuals with anomia and in individuals with Wernicke’s aphasia using a task similar to that used in Lupyan and Mirman (2013). Others described aphasia-related categorization deficits for both HD and LD categories (Koemeda-Lutz et al., 1987) or no deficits in either (Hough, 1993). Further, variations in the task (such as showing the category label to the participant during the entire trial vs. just at the beginning of the trial) significantly affected categorization performance in participants with aphasia (Koemeda-Lutz et al., 1987), suggesting that task demands may contribute to the observed results (above and beyond alleged effects of category type). Finally, some have argued for a relationship between categorization difficulties and conceptual-semantic rather than linguistic impairments (Caramazza et al., 1982; Whitehouse et al., 1978; cf. Le Dorze & Nespoulous, 1989).

It is important to emphasize that even if individuals with aphasia did consistently show a selective impairment in LD categorization, it would not necessarily implicate language as the source of the deficit. In particular, the language network in the left hemisphere, especially in the left frontal cortex, lies adjacent to the domain-general multiple demand network, which supports executive functions, like working memory and inhibitory control (Assem, Glasser, et al., 2020; Duncan, 2010, 2013; Fedorenko et al., 2012, 2013). As a result, left hemisphere damage can lead to joint linguistic and domain-general executive deficits (Baldo et al., 2010; Gainotti et al., 1986). Prior work has shown that performance on executive function tasks, not language tasks, predicts success in learning novel categories (Vallila-Rohter & Kiran, 2015). Further, the multiple demand network, but not the language network, is robustly sensitive to cognitive effort across domains (e.g., Fedorenko et al., 2011, 2013; Hugdahl et al., 2015; Shashidhara et al., 2019), and LD categorization appears to be more cognitively challenging than HD categorization: LD categories are harder to learn for both human children (e.g., Kloos & Sloutsky, 2008) and non-human primates (Couchman et al., 2010), require supervision (e.g., Kloos & Sloutsky, 2008), and are generally linked with executively-taxing intentional learning (Ashby et al., 1998; Ashby & Ell, 2001; Ashby & O’Brien, 2005; Couchman et al., 2010; Kemler Nelson, 1984). Finally, information about LD category membership is typically not stored but rather “computed on the fly”, which can also result in higher cognitive load. It is therefore possible that impaired performance on LD categorization (and on categorization tasks more broadly) depends primarily on domain-general executive resources.

The second prediction of the LD-specific language recruitment hypothesis is that LD categories would evoke stronger activity within the language brain regions. To our knowledge, this hypothesis has not been directly tested in the neuroimaging literature; instead, many studies have investigated differences between taxonomic and thematic relations (e.g., Kalénine et al., 2009; Lewis et al., 2015; Sachs et al., 2008; Sass et al., 2009), both of which are considered HD. Further, few neuroimaging studies employ methods that would be required to dissociate the contributions of language-specific regions from those of domain-general executive regions: given the inter-individual variability in the precise locations of functional areas, voxels in anatomically identical locations within the frontal lobe might be language-specific in one individual and domain-general in another, so traditional group-based analyses (Friston et al., 1994) would fail to distinguish between them (Fedorenko et al., 2012; Fedorenko & Blank, 2020; Nieto-Castañón & Fedorenko, 2012). Unambiguously assessing the role of language in LD categorization requires identification of language-specific regions in individual participants and testing their responses to LD compared to HD conditions.

### 1.3 Current study

Here, we report three interlinked experiments aimed at re-examining the role of language in LD and HD categorization. In line with recent emphasis on robust and replicable science (e.g., Ioannidis, 2014; Poldrack et al., 2017), in Experiments 1 and 2, we attempt to conceptually replicate the findings of Lupyan and Mirman (2013; L&M henceforth). In Experiment 1, we closely follow L&M’s experimental procedure, but additionally include another brain-damaged control group (individuals with Parkinson’s disease, or PD) to examine general effects of brain damage on performance. In Experiment 2, we adjust the experimental paradigm to reduce general executive demands, which might affect performance (e.g., Koemeda-Lutz et al., 1987). Finally, in Experiment 3, we use fMRI in neurotypical individuals to test the prediction that the language system is engaged during LD categorization more than during HD categorization.

To foreshadow our results, we find that participants with aphasia perform worse on the categorization task overall, but this effect does not consistently and selectively affect LD categories. In Experiment 1, participants with aphasia actually performed better on LD trials than on HD trials. In Experiment 2, participants with low naming scores did show impaired performance on LD categorization compared to HD categorization; however, the performance of one individual with severe naming impairments was within 2 standard deviations of healthy controls on both LD and HD categorization. Finally, Experiment 3 revealed low engagement of the language network during both LD and HD categorization, with no significant difference between the two. Thus, the influence of language on LD categorization is behaviorally inconsistent and is not supported by fMRI evidence, leading us to conclude that the language system does not play a special role in LD (single-feature-based) categorization and is not engaged during categorization in general.

## 2. Experiment 1

The aim of Experiment 1 was to replicate the effects reported by L&M by using a closely related experimental paradigm. In their study, L&M compared LD and HD categorization performance in participants with anomic aphasia and in neurotypical controls. They found a) lower performance on LD compared to HD categories in both healthy adults and participants with anomic aphasia; and, critically, b) a greater decrement in performance for the LD, compared to the HD condition in participants with aphasia. We explored whether these same effects would be present in our study. To additionally examine the extent to which performance might depend on the general effect of brain damage, as opposed to a linguistic impairment, we also included a group of individuals with Parkinson’s disease (PD).

### 2.1 Methods

#### 2.1.1 Participants

Neurotypical older participants (*N*=9 (6 F), age *M*=67.89, *SD*=14.98) were recruited by convenience sampling; individuals with chronic aphasia (*N*=11 (3 F), age *M*=61.18, *SD*=12.09) were recruited from the UCL Aphasia Clinic Research Register. The aphasia group included patients with a range of aphasia types and severities. Unlike L&M, we did not try to limit our sample to individuals classified as having “Anomic” aphasia, given that the use of such rigid classification labels fails to account for the heterogeneity among the symptoms observed across patients (Badecker & Caramazza, 1985; Caramazza & Badecker, 1989), and given that some degree of anomia is present in all forms of aphasia (e.g., Blumstein, 1988; Goodglass & Geschwind, 1976). Instead, we explicitly estimated the patients’ performance on tasks of interest (see below). Individuals with PD (*N*=14 (8 F), age *M*=68.64, *SD*=11.69) were recruited from the Parkinson’s UK Research Registry. For detailed participant information, see **Table 1**. All participants used English as their primary language. Patients were offered a £10.00 reimbursement. Ethical approval was granted by the UCL Research Ethics panel, Project ID: LC/2013/05, and all volunteers gave informed consent to participate in the experiment.

**Table 1.** Participant information, Experiment 1.

| Group | Participant | Age | Education | Gender | TPO<br>(months) | BNT |
| --- | --- | --- | --- | --- | --- | --- |
| Neurotypical | 1 | 75 | Up to 16 | F | - | 51 |
|  | 2 | 68 | Up to 16 | F | - | 55 |
|  | 3 | 68 | Up to 16 | M | - | 55 |
|  | 4 | 56 | Degree-Level | F | - | 59 |
|  | 5 | 98 | Up to 16 | F | - | 47 |
|  | 6 | 54 | Degree-Level | M | - | 53 |
|  | 7 | 69 | Up to 16 | M | - | 55 |
|  | 8 | 76 | Up to 16 | F | - | 52 |
|  | 9 | 47 | Up to 18 | F | - | 58 |
| PD | 1 | 60 | Postgraduate | M | 36 | 59 |
|  | 2 | 58 | Degree-Level | M | 12 | 58 |
|  | 3 | 80 | Up to 18 | F | 48 | 58 |
|  | 4 | 56 | Postgraduate | F | 48 | 54 |
|  | 5 | 66 | Degree-Level | F | 72 | 59 |
|  | 6 | 75 | Degree-Level | F | 96 | 56 |
|  | 7 | 76 | Up to 18 | M | 36 | 54 |
|  | 8 | 59 | Degree-Level | F | 60 | 55 |
|  | 9 | 69 | Postgraduate | F | 36 | 54 |
|  | 10 | 63 | Postgraduate | F | 60 | 56 |
|  | 11 | 77 | Degree-Level | M | 12 | 46 |
|  | 12 | 72 | Postgraduate | M | 120 | 53 |
|  | 13 | 75 | Degree-Level | M | 2 | 58 |
|  | 14 | 75 | Postgraduate | F | 360 | 53 |
| Aphasia | 1 | 52 | Degree-Level | M | 120 | 30 |
|  | 2 | 57 | Up to 16 | M | 84 | 57 |
|  | 3 | 52 | Up to 18 | M | 48 | 52 |
|  | 4 | 59 | Postgraduate | M | 120 | 43 |
|  | 5 | 79 | Up to 16 | F | 36 | 50 |
|  | 6 | 44 | Up to 18 | F | 12 | 14 |
|  | 7 | 81 | Up to 16 | M | 96 | 57 |
|  | 8 | 56 | Up to 18 | M | 60 | 12 |
|  | 9 | 57 | Up to 18 | M | 48 | 51 |
|  | 10 | 60 | Up to 16 | M | 132 | 34 |
|  | 11 | 76 | Up to 16 | F | 84 | 14 |

TPO- Time Post Onset; BNT- Boston Naming Test

#### 2.1.2 Design and Materials

The critical categorization task was modeled closely on L&M’s experiment, which used 34 unique categories (18 HD categories and 16 LD categories), with some repetition of categories in each condition. We chose to not repeat any categories, so we limited the materials to 16 categories in each condition (dropping ‘BODY PARTS’ and ‘FACIAL FEATURES’ from the HD set). Unlike L&M, who used normed color drawings (Rossion & Pourtois, 2004), we used high-quality color photographs selected from the Hemera Photo Objects 5000 and Google Images. For each category, we selected 8-15 targets and 25-27 distractors. Distractors included some items which were related to the target category (for example, for the category ‘DANGEROUS ANIMALS’, 13 of the 26 distractors were animals that were not dangerous, and the category ‘ANIMALS WITH STRIPES’ included distractors that were animals without stripes, and inanimate objects with stripes). A total of 1,087 unique images were used (any given image appeared as a target in 0-2 categories and as a distractor in 0-2 categories). All photographs depicted objects on a white background. The materials and the experimental scripts for all experiments are available on OSF: https://osf.io/guwh8/.

To determine the extent of lexical impairment in the aphasia group and to compare lexical abilities across the three groups, all participants completed the Boston Naming Test (BNT; Goodglass et al., 1983), where they were sequentially presented with up to 60 line drawings of objects and asked to overtly name each one. The standard discontinuation rule was applied, with testing stopped after eight consecutive failed naming attempts. No semantic or phonological cues were given.

#### 2.1.3 Experimental Procedure

Testing was carried out individually either in a quiet well-lit room at the UCL Aphasia clinic or at the participants’ home, using a MacBook Pro (Retina, 13-inch display) and an external computer mouse. The experiment was set up using PsychoPy (Version 1.83), and the procedure closely followed that used in L&M’s experiment, except where noted. On each trial (see **Figure 1** **(top)** for a sample HD and LD trial), participants were presented with a 4 x 5 grid of images. The image sets for the individual trials—each consisting of 20 images (4 targets and 16 distractors)—were randomly selected from the pool of targets/distractors for each participant separately. The category was stated at the top of the screen in lower-case Arial bold letters (e.g., ‘objects that hold water’) and remained on the screen for the duration of the trial. Participants selected the objects that belonged to the target category by clicking on each relevant image. A gray frame appeared around an image once it was clicked; clicking the image again de-selected it (removed the gray frame) to allow participants to modify responses. Once the participant had selected all of the images they deemed appropriate for the target category, they clicked a large green button with the word ‘Done’ at the bottom of the screen (in the L&M version, the button said ‘click here when done’). Doing so triggered the next trial. Although each trial contained a fixed number of targets (four), participants were not informed of the number of targets during the instructions and could therefore select as many images as they wished on any given trial. No time limit was imposed on the trials, but participants were encouraged to work as quickly and accurately as possible. HD and LD trials were interleaved, and the order of conditions was randomized for each participant. Each participant performed the experiment twice for a total of 64 trials (32 per condition), but in contrast to L&M, different sets of images were used for the two instances of each category to minimize practice effects. Responses were recorded for each image; response times were recorded for each trial (the time from the onset of the trial until the ‘Done’ button was pressed^3^). The experiment lasted approximately one hour. The BNT (Goodglass et al., 2001) was administered between the two runs of the experiment.

#### 2.1.4 Statistical analyses

To determine possible differences in demographics and BNT scores across groups, we conducted ANOVA tests (with follow-up Bonferroni-corrected t-tests), implemented in SPSS 22 (IBM Corp., 2013). For the critical analyses, we used linear/logistic mixed effect regression models (Baayen et al., 2008). Given that correct or incorrect selection of items is categorical in nature, we use logistic regression to analyze accuracy measures (Jaeger, 2008). For response times, we use linear regression. When specifying model contrasts, we used sum coding for category dimension (HD vs. LD); the effect of group was therefore estimated across both category dimensions. For participant group (neurotypical vs. aphasia vs. PD), we used dummy coding with ‘neurotypical’ as the reference level; thus, the effect of category was estimated specifically for the neurotypical group (with interaction terms denoting whether the category effect differed for the aphasia/PD groups). For completeness and to facilitate result comparison with L&M, we also ran pairwise comparisons across groups using ‘aphasia’ as the reference level (the results were Bonferroni-corrected, n=2). The mixed effect analyses were run using the *lmer* function from the *lme4* R package (D. Bates et al., 2015); statistical significance of the effects was evaluated using the *lmerTest* package (Kuznetsova et al., 2017); follow-up comparisons were conducted using the *emmeans* package (https://cran.r-project.org/package=emmeans). Lastly, due to a technical error, if participants accidently double-clicked the ‘Done’ button, the next set of images was skipped, and the software registered it as though no response was made by participants. As a result, we excluded trials where no selection was made and where the trial length was less than 5 seconds. This resulted in the exclusion of 42 trials (out of 2177), spread randomly between participants, groups and categories. The analysis code is available on OSF: https://osf.io/guwh8/.

### 2.2 Results

#### 2.2.1 Group profiles

As expected, the neurotypical, aphasia, and PD groups differed significantly in their BNT scores (*F*(2,31)=9.85, *p*<.001). Post-hoc pairwise comparisons showed that the BNT scores of participants with aphasia (*M*=37.64, *SD*=17.78) were significantly lower than those of neurotypical participants (*p*=.005) or participants with PD (*p*=.001), with the latter two groups not differing significantly (*M*=53.89, *SD*=3.66 vs. *M*=55.21, *SD*=3.42; *p*>.999). The groups did not differ in age (*F*(2,31)=1.45, *p*=.250), but a significant difference was observed in the level of education (*F*(2,31)=14.36, *p*<.001): participants with PD group were significantly more educated than both neurotypical participants (*p*=.001) and participants with aphasia (*p* =.002), with the latter two not differing significantly (*p*>.999).

#### 2.2.2 Categorization task

Following L&M, we analyzed three dependent variables: hit rate (the number of targets selected out of 4), false alarm rate (the number of distractors selected out of 16), and trial response time (RT). To account for heterogeneity among participants and experimental categories, we used mixed effect regression models (Baayen et al., 2008) with category dimension (HD vs. LD), group (neurotypical, aphasia, and PD) and their interaction as fixed effects, as well as category (e.g. “DANGEROUS ANIMALS”) and participant ID as random effect intercepts. Categorization results are summarized in **Figure 2**.

**Figure 2.**
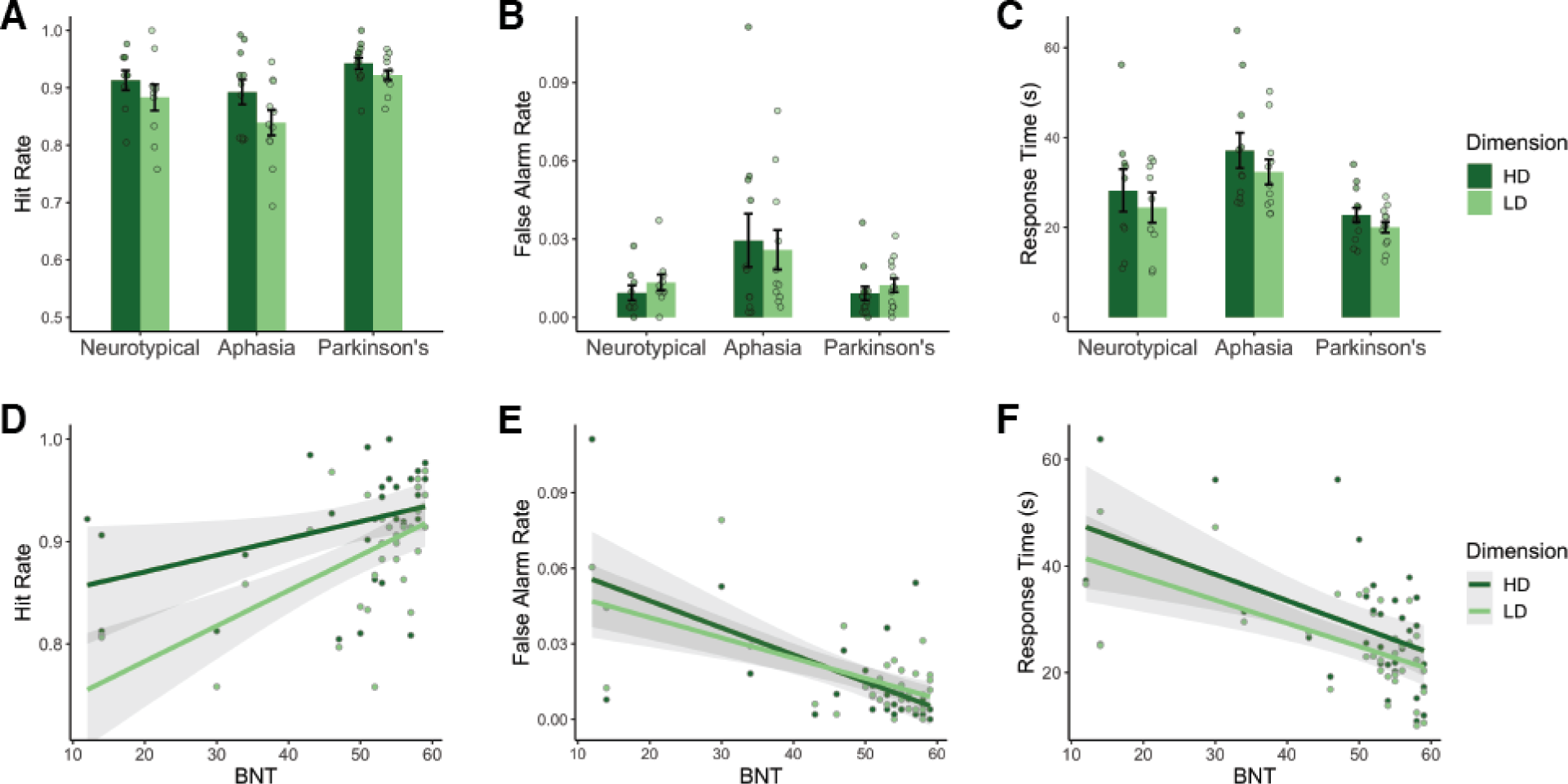
Experiment 1 results. (**A**) Hit Rate, (**B**) False Alarm Rate, and (**C**) Response Time (RT) across the three participant groups (here, RT is the time from trial onset until participants pressed the “Done” button). (**D)** Hit Rate, (**E**) False Alarm rate, and (**F**) RT plotted against participants’ BNT scores, a measure of naming performance. Here and elsewhere, error bars depict the standard error across participants.

##### Hit rate

Participants with aphasia had similar hit rates for LD categories (*M*=0.84, *SD*=0.07) and HD categories (*M*=0.89, *SD*=0.07; LD>HD: *β*=-0.41, *SE*=0.28, *p*=.139). The overall hit rate for participants with aphasia (*M*=0.87, *SD*=0.08) was similar to neurotypical participants (*M*=0.90, *SD*=0.06; neurotypical>aphasia: *β*=0.24, *SE*=0.25, *p*=.338) and lower than for participants with PD (*M*=0.93, *SD*=0.03; PD>aphasia: *β*=0.72, *SE*=0.23, *p*=.002). Moreover, we did not observe a reliable category dimension by group interaction for the aphasia vs. neurotypical comparison (*β*=0.04, *SE*=0.18, *p*=.813), nor for the aphasia vs. PD comparison (*β*=0.19, *SE*=0.18, *p*=.304). Follow-up analyses showed that there was no main effect of category dimension across groups (*β*=0.34, *SE*=0.27, *p*=.813), nor within the neurotypical group (*β*=0.37, *SE*=0.29, *p*=.478) or the PD group (*β*=0.22, *SE*=0.29, *p*=.788). These results fail to replicate the findings by L&M, who reported the main effect of category dimension, as well as a selective impairment in LD categorization for patients with aphasia (**Table 2**).

**Table 2.**
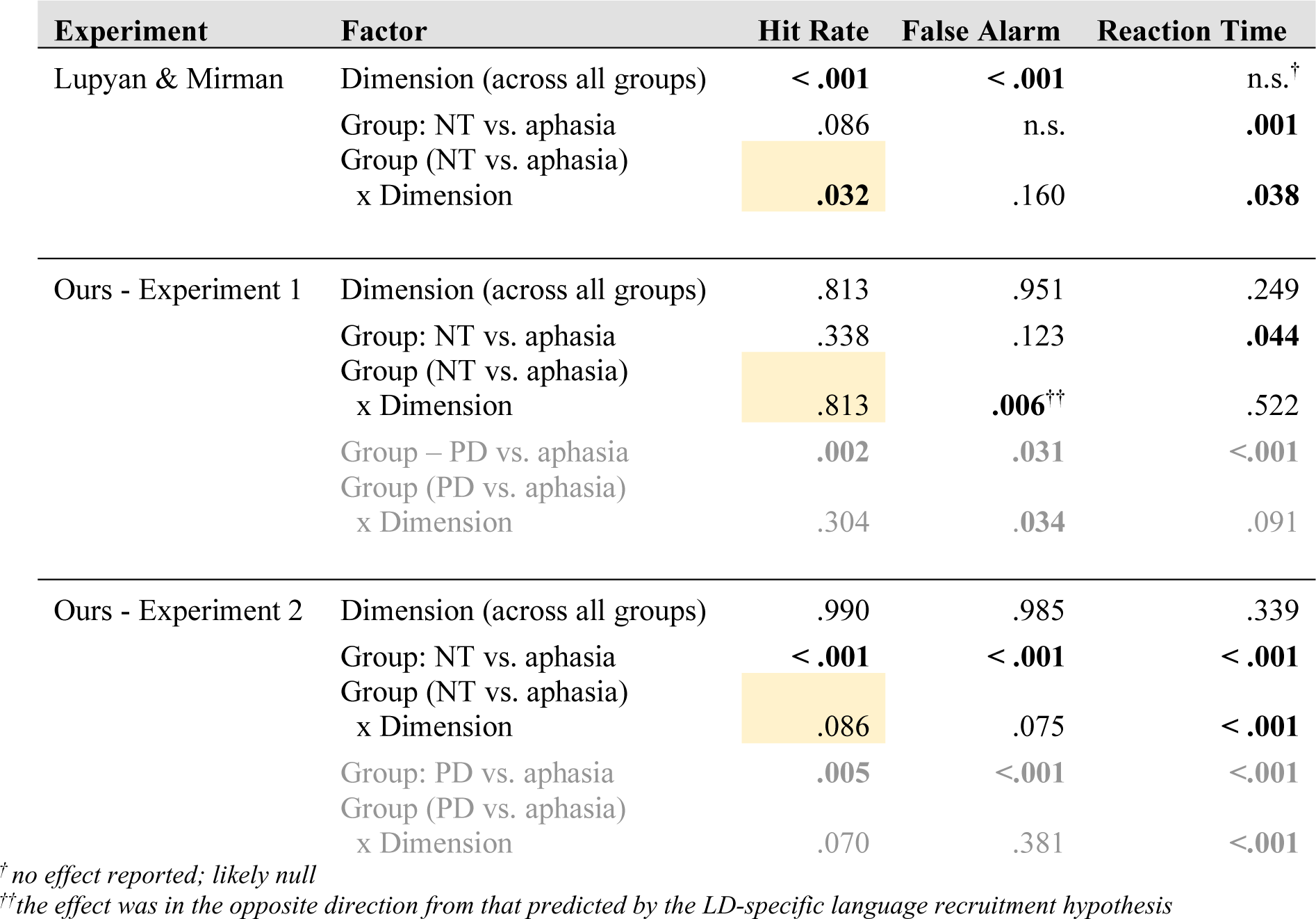
Statistical significance (p values) of group and category dimension effects on categorization performance. The critical result reported by L&M is the Aphasia/Neurotypical (NT) Group by LD/HD interaction — highlighted in yellow — that we do not replicate. Here and below, p values in the L&M paper were obtained with ANOVA tests, whereas p values in our experiments were obtained with linear/logistic mixed effect models. See the main text for analysis details.

We additionally conducted an exploratory analysis to investigate the difference between the aphasia and PD groups. Given that the PD group had a higher average education level, we repeated the analysis above with ‘education level’ as an additional fixed effect. The updated model had a similar fit to the data compared to the original (as per the likelihood ratio test: χ^2^=3.39, *p*=.065); under this model, the difference between the aphasia and the PD groups was no longer significant (*β*=0.37, *SE*=0.29, *p*=.202). The significance of other effects was unchanged.

##### False alarm rate

The false alarm rate in participants with aphasia also did not differ between LD categories (*M*=0.03, *SD*=0.03) and HD categories (*M*=0.03, *SD*=0.03; LD>HD: *β*=-0.22, *SE*=0.35, *p*=.534). As with the hit rate, the overall false alarm rate for participants with aphasia (*M*=0.03, *SD*=0.03) was comparable to that of neurotypical participants (*M*=0.01, *SD*=0.01; neurotypical>aphasia: *β*=-0.58, *SE*=0.37, *p*=.123), although participants with PD performed better than participants with aphasia, i.e., with fewer false alarms (*M*=0.01, *SD*=0.01; PD>aphasia: *β*=-0.74, *SE*=.34, *p*=.031). Unlike the hit rate results above, there was a significant interaction between category dimension (LD>HD) and group (neurotypical>aphasia: *β*=0.63, *SE*=0.23, *p*=.006; PD>aphasia: *β*=0.44, *SE*=0.21, *p*=.034). However, this interaction effect goes in the opposite direction from that predicted by the LD-specific language recruitment hypothesis: participants with aphasia performed *better* on LD categories relative to controls. The pattern of results is also inconsistent with L&M’s results in that they found no interaction between group and category dimension. Lastly, follow-up analyses showed no main effect of category dimension across groups (*β*=-0.14, *SE*=0.34, *p*=.951), nor within the neurotypical group (*β*=-0.41, *SE*=0.38, *p*=.614) or the PD group (*β*=-0.22, *SE*=0.37, *p*=.879).

Similar to the hit rate analysis, an exploratory model that included ‘education level’ as a fixed effect explained a similar amount of variance compared to the original model (χ^2^=0.35, *p*=.557) and no longer showed a significant difference between the aphasia and PD groups (β=-0.57, *SE*=0.45, *p*=.209). The significance of other effects was unchanged.

##### Response time

The RT analysis revealed that participants with aphasia were faster to respond during LD trials (*M*=32.36, *SD*=9.33) compared to HD trials (*M*=37.10, *SD*=12.90; LD>HD: *β*=-4.75, *SE*=2.26, *p*=.042), in contrast to the predictions of the LD-specific language recruitment hypothesis. The overall RTs for participants with aphasia (*M*=34.70, *SD*=11.30) were longer than for neurotypical participants (*M*=26.30, *SD*=12.10; *β*=-8.42, *SE*=4.02, *p*=.044) and the PD group (*M*=21.40, *SD*=5.14; *β*=-13.30, *SE*=3.66, *p*<.001). The interactions between group and category dimension were not significant (neurotypical>aphasia: *β*=0.82, *SE*=1.29, *p*=.522; PD>aphasia: *β*=1.98, *SE*=1.17, *p*=.091). Follow-up analyses showed no overall effect of category dimension across groups (*β*=3.81, *SE*=2.19, *p*=.249), within the neurotypical group (*β*=3.92, *SE*=2.34, *p*=.271) or within the PD group (*β*=2.76, *SE*=2.27, *p*=.521).

##### Effect of naming performance

To explore the effect of naming ability on the categorization task performance, we fitted a logistic mixed effect linear regression model with the BNT score, category dimension, and their interaction as fixed effects and participants (across the three groups) and categories (e.g., “DANGEROUS ANIMALS”) as random effects. Similar to L&M, we also included education level as a fixed effect.

We found that BNT was a significant predictor of hit rate (*β*=0.28, *SE*=0.08, *p*<.001), false alarm rate (*β*=-0.47, *SE*=0.12, *p*<.001), and RT (*β*=-5.26, *SE*=1.41, *p*<.001), such that higher BNT scores corresponded to more accurate and faster performance (**Figure 2D-F**). There was no main effect of category dimension (hit rate: *β*=-0.31, *SE*=0.27, *p*=.236; false alarm rate: *β*=0.14, *SE*=0.35, *p*=.690; RT: *β*=-3.73, *SE*=2.14, *p*=.092). However, we observed an interaction between BNT and category dimension for false alarm rate (*β*=0.16, *SE*=0.16, *p*=.011), such that participants with lower BNT scores had higher false alarm rates for HD compared to LD categories. No interaction was observed for group and category dimension for hit rate (*β*=0.11, *SE*=0.07, *p*=.078) or RT (*β*=0.74, *SE*=0.49, *p*=.131). Conversely, education was a significant predictor for hit rate (*β*=0.23, *SE*=0.07, *p*=.001) and RT (*β*=-2.95, *SE*=1.24, *p*=.024), but not for false alarm rate (*β*=-0.09, *SE*=0.11, *p*=.436). These results do not replicate the pattern of results reported by L&M (**Table 3**) and do not support the LD-specific language recruitment hypothesis.

**Table 3.** Statistical significance (p values) of category dimension and anomia effects on categorization performance. The critical result reported by L&M is the Aphasia/Neurotypical Group by LD/HD interaction — highlighted in yellow — that we replicate in Experiment 2 but not in Experiment 1. The effect of naming performance on response times is not reported in L&M, so we omit it from this table.

| Experiment | Factor | Hit Rate | False Alarm |
| --- | --- | --- | --- |
| Lupyan & Mirman | Dimension | ?<sup>†</sup> | ?<sup>†</sup> |
|  | PNT | **.015** | n.s. |
|  | PNT x Dimension | **.015** | n.s. |
| Ours - Experiment 1 | Dimension | .236 | .690 |
|  | BNT | **<.001** | **<.001** |
|  | BNT x Dimension | **.078** | **.011<sup>††</sup>** |
| Ours - Experiment 2 | Dimension | .885 | .639 |
|  | BNT | **.002** | **<.001** |
|  | BNT x Dimension | **.039** | .090 |

PNT – Philadelphia Naming Test; BNT – Boston Naming Test
<sup>†</sup> no effect reported
<sup>††</sup> the effect was in the opposite direction from that predicted by the LD-specific language recruitment hypothesis

### 2.3 Interim Discussion

In Experiment 1, we tested the hypothesis that language is selectively recruited to support LD categorization by using a setup similar to L&M’s study. Our findings differed from those reported in the original paper. L&M reported that (a) performance was better for HD than for LD categories, (b) hit rate but not false alarm rate was predicted by naming ability, and (c) individuals with aphasia were selectively impaired on LD categorization, as indicated by hit rate (but not false alarm rate). In our results, none of the three outcome measures (hit rate, false alarm rate, RT) differed between LD and HD categorization conditions, both hit rate and false alarm rate were predicted by naming ability, and there was no interaction between group and dimension for hit rate. Moreover, we observed a significant interaction between group and dimension for the false alarm rate that went in the opposite direction from that predicted by the original hypothesis, i.e., participants with aphasia performed better on LD categories relative to controls. In summary, Experiment 1 provides no support for the hypothesis that language plays a special role in LD categorization.

With regard to group differences, participants with aphasia performed as accurately as the neurotypical controls. Participants with PD performed better, but this difference is likely explained by the higher education level reported by participants in this group. As in L&M, our findings showed that participants with aphasia were significantly slower to complete the categorization task compared to the neurotypical and PD groups. However, the reason for this slower performance can be explained by the presence of more severe motor impairments in participants with aphasia than participants with PD (e.g., right hemiplegia), often necessitating use of their non-preferred hand. Thus, we are hesitant to place a lot of weight on the RT differences and primarily focus on the hit rate and false alarm results.

Across groups, BNT scores significantly predicted performance on all three outcome measures (although this effect did not differ for LD and HD categorization). This relationship has at least two possible explanations. First, the categorization task, as designed by L&M, does require some linguistic processing: the participants need to read and understand the label, which often consists of multiple words (e.g., “NON_FOOD THINGS FOUND IN THE KITCHEN”). Thus, a disruption in receptive language may make the categorization task more difficult for individuals with aphasia. Under this explanation, BNT scores may be a proxy for overall aphasia severity. The second explanation is that, due to the proximity of language-specific and multiple-demand brain regions in some parts of the brain (Fedorenko et al., 2012; Fedorenko & Blank, 2020), brain damage that causes lower BNT scores also leads to difficulties with executively challenging tasks. The categorization task adopted from L&M involves visual search and selecting among multiple options, which require substantial executive involvement (Petersen & Posner, 2012; Posner & Petersen, 1990); thus, categorization difficulties might reflect this increased recruitment of executive demand resources.

Why did we fail to find support for the LD-specific language recruitment hypothesis? One possibility is that the effect is not stable. The findings from L&M did not show large effect sizes, and given the number of participants in neuropsychological studies is typically not large, this could result in effects that are volatile. Another explanation is that the selective impairment in LD categorization manifests only in individuals with very low naming performance. Thus, we might have missed the effect of interest because we recruited participants with a fairly wide range of naming scores. To address these potential concerns, we conducted a modified version of the categorization experiment with a new set of participants, with a focus on individuals with severe naming impairments in the aphasia group.

## 3. Experiment 2

The aim of Experiment 2 was three-fold. First, we wanted to follow up on the relationship between naming ability (BNT scores) and categorization performance, which was reported by L&M and found in Experiment 1. Thus, we recruited participants with aphasia who had severe anomia, as measured by the BNT (score range 1-11, compared to 12-57 in Experiment 1; see **Tables 1 and 4**). Second, we adjusted the paradigm to minimize executive demands, including attention, visual search, selection/inhibition, and updating. Third, we sought to validate a version of the task that could be used in an fMRI setting (time-locked to events). See **Figure 1** **(bottom)** for the modified task setup.

**Table 4.** Participant information, Experiment 2.

| Group | Participant | Age | Education | Gender | TPO<br>(months) | BNT |
| --- | --- | --- | --- | --- | --- | --- |
| Neurotypical | 1 | 68 | Degree-Level | F | - | 51 |
|  | 2 | 61 | Postgraduate | F | - | 41 |
|  | 3 | 85 | Degree-Level | F | - | 54 |
|  | 4 | 73 | Postgraduate | F | - | 58 |
|  | 5 | 72 | Up to 18 | F | - | 58 |
|  | 6 | 77 | Postgraduate | F | - | 55 |
|  | 7 | 77 | Degree-Level | F | - | 59 |
|  | 8 | 66 | Degree-Level | F | - | 57 |
|  | 9 | 66 | Postgraduate | F | - | 54 |
|  | 10 | 76 | Degree-Level | F | - | 59 |
|  | 11 | 65 | Postgraduate | F | - | 56 |
|  | 12 | 80 | Up to 18 | F | - | 45 |
|  | 13 | 74 | Postgraduate | F | - | 54 |
|  | 14 | 71 | Degree-Level | F | - | 57 |
|  | 15 | 76 | Degree-Level | F | - | 47 |
| PD | 1 | 71 | Degree-Level | M | 24 | 58 |
|  | 2 | 78 | Degree-Level | M | 24 | 47 |
|  | 3 | 64 | Postgraduate | M | 30 | 48 |
|  | 4 | 72 | Postgraduate | M | 18 | 59 |
|  | 5 | 54 | Degree-Level | M | 204 | 58 |
|  | 6 | 72 | Degree-Level | M | 4 | 48 |
|  | 7 | 62 | Postgraduate | F | 120 | 56 |
|  | 8 | 65 | Postgraduate | M | 17 | 59 |
|  | 9 | 74 | Up to 18 | M | 96 | 56 |
|  | 10 | 67 | Up to 16 | M | 60 | 54 |
|  | 11 | 67 | Postgraduate | M | 72 | 59 |
|  | 12 | 59 | Postgraduate | M | 30 | 58 |
|  | 13 | 59 | Degree-Level | M | 48 | 60 |
|  | 14 | 67 | Degree-Level | M | 18 | 55 |
|  | 15 | 68 | Degree-Level | M | 98 | 48 |
| Aphasia | 1 | 58 | Up to 18 | M | 42 | 5 |
|  | 2 | 68 | Up to 16 | M | 68 | 9 |
|  | 3 | 77 | Up to 18 | M | 111 | 11 |
|  | 4 | 57 | Degree-Level | M | 34 | 1 |
|  | 5 | 73 | Up to 18 | M | 326 | 4 |

TPO- Time Post Onset; BNT- Boston Naming Test

### 3.1 Method

#### 3.1.1 Participants

Neurotypical participants (*N*=15 (15 F), age *M*=72.47, *SD*=6.41) were recruited by convenience sampling; patients with chronic aphasia and severe lexical access impairment (*N*=5 (all males), age *M*=66.60, *SD*=8.91) were recruited from Aphasia volunteer research registers; PD patients (*N*=15 (1 F), age *M*=66.60, *SD*=6.38) were recruited from the Parkinson’s UK Research Registry (see **Table 4** for detailed participant information). None of the participants took part in Experiment 1. All participants used English as their primary language and were offered a £15.00 reimbursement. Ethical approval was granted by the UCL Research Ethics panel, Project ID: LC/2013/05, and all volunteers gave informed consent to participate in the experiment.

#### 3.1.2 Design and Materials

The categories were identical to those of Experiment 1. The images were also largely the same although some were replaced by better quality photographs. Unlike Experiment 1, we presented the images sequentially (**Figure 2, bottom**). Each block started with a category label, followed by 12 images presented one at a time. The category label remained on the screen to minimize memory demands. The images for each category block were randomly selected from the general set of pictures for that category. The number of targets varied across blocks (minimum: 4, maximum: 6) so as to minimize the implicit learning of a fixed number of targets, which could have incentivized participants to keep track of the total number of targets and thereby increase their cognitive load. Categories were grouped by dimension (LD/HD) into groups of 4, for a total of 8 blocks (4 blocks per dimension). These 8-block sequences (“runs”) were separated by a rest period of fixation (10s in duration). The order of runs, the order of conditions within runs (LD first vs. HD first), the order of categories within runs, and the order of images within category blocks were randomized for each participant.

#### 3.1.3 Experimental Procedure

Testing was carried out individually either in a quiet well-lit room at a clinic nearest to the participant’s location or in their home, using a Dell Latitude E5540 (14.1-inch display). The experiment was set up using Python (version 2.7.10). Each category block started with an instruction screen presented for 2s that read ‘Please find [CATEGORY LABEL]’ (e.g., ‘Please find objects that hold water’). Given that the participants in the aphasia group were severely lexically impaired and had difficulty processing orthographic information, the experimenter read the category label aloud to all participants (in all groups) during this trial-initial 2s window. This screen was followed by a sequence of 12 images presented one at a time for a maximum of 10s per image. For each image, participants had to decide whether the depicted object belong to the target category by pressing one of two keys on the keyboard: the ‘Y’ key marked with a green sticker for YES, or the ‘N’ key marked with a red sticker for NO. If no response was recorded for 10s, the experiment advanced to the next image. Responses and response times were recorded for each image. The experiment lasted approximately 1 hour. The BNT was administered at the beginning of the testing session.

#### 3.1.4 Statistical analyses

The statistical analysis procedure was the same as in Experiment 1. No trials were excluded.

### 3.2 Results

#### 3.2.1 Group profiles

As expected, the groups differed significantly in their BNT scores (*F*(2,32)=202.67, *p*<.001). Post-hoc pairwise comparisons revealed that the BNT scores of participants with aphasia (*M*=6.00, *SD*=4.00) were significantly lower than both neurotypical participants (*p*<.001) and participants with PD (*p*<.001), with the latter two groups not differing significantly (*M*=53.67, *SD*=5.42 vs. *M*=54.87, *SD*=4.73, *p>*.999). The groups did not differ in age (*F*(2,32)=3.23, *p*=.053), but a significant difference was observed in the level of education (*F*(2,32)=5.42, *p*=.009), with neurotypical participants and participants with PD having significantly more years of education than participants with aphasia (*p=*.010 and *p=*.016, respectively). The neurotypical participants and participants with PD did not differ (*p* >.999).

#### 3.2.2 Categorization task

As in Experiment 1, three dependent variables were analyzed: hit rate, false alarm rate, and RT. Categorization results are summarized in **Figure 3**.

**Figure 3.**
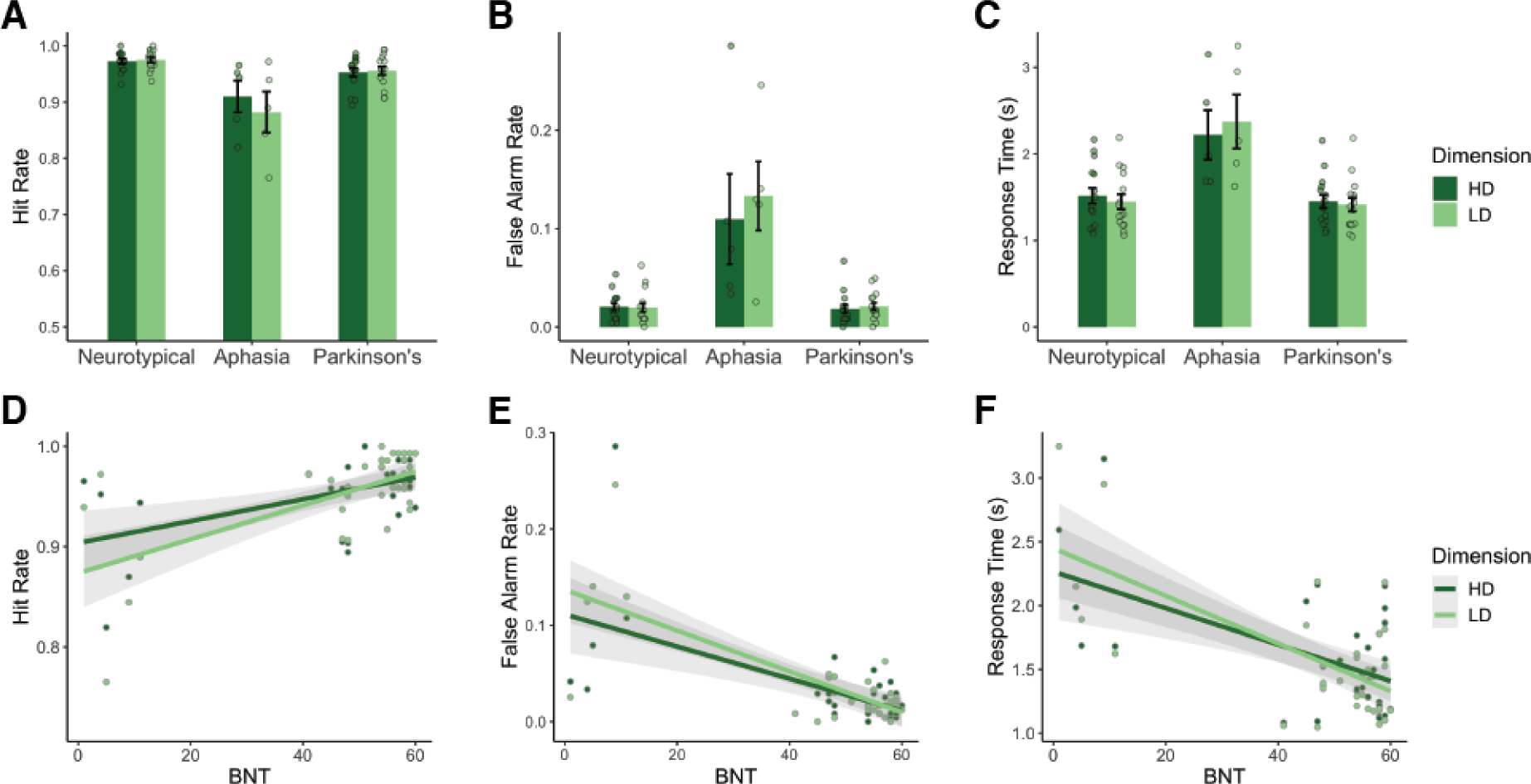
Experiment 2 results. (**A**) Hit Rate, (**B**) False Alarm Rate, and (C) Response Time (RT) across the three participant groups (here, RT is the time until participants pressed a “yes” or “no” button for each image within a trial). (**D**) Hit Rate, (**E**) False Alarm Rate, and (**F**) RT plotted against participants’ BNT scores, a measure of naming performance.

#### Hit rate

Similar to the results of Experiment 1, participants with aphasia had similar hit rates for LD categories (*M*=0.88, *SD*=0.08) and HD categories (*M*=0.91, *SD*=0.06; LD>HD: *β*=-0.34, *SE*=0.29, *p*=.252). Participants with aphasia had overall lower hit rates (*M*=0.90, *SD*=0.07) compared to neurotypical participants (*M*=0.97, *SD*=0.02; neurotypical>aphasia: *β*=1.45, *SE*=0.32, *p*<.001) and participants with PD (*M*=0.95, *SD*=0.03; PD>aphasia: *β*=0.86, *SE*=0.31, *p*=.005), which is consistent with Experiment 1’s negative relationship between naming ability and categorization performance. We did not observe a reliable category dimension by group interaction for the aphasia vs. neurotypical comparison (*β*=0.44, *SE*=0.26, *p*=.086), nor for the aphasia vs. PD comparison (*β*=0.42, *SE*=0.23, *p*=.070). Follow-up analysis showed that there was no main effect of category dimension across groups (*β*=0.05, *SE*=0.25, *p*=.990), nor within the neurotypical group (*β*=-0.11, *SE*=0.30, *p*=.960), or the PD group (*β*=-0.08, *SE*=0.28, *p*=.975). Overall, the group comparison of hit rate (**Figure 3A**) does not support the LD-specific language recruitment hypothesis.

#### False alarm rate

The false alarm rate results (**Figure 3B**) were consistent with the hit rate results. Participants with aphasia had comparable false alarm rates for LD categories (*M*=0.13, *SD*=0.08) and HD categories (*M*=0.11, *SD*=0.10; LD>HD: *β*=0.13, *SE*=0.27, *p*=.626). The overall false alarm rate among participants with aphasia (*M*=0.12, *SD*=0.09) was higher than in neurotypical participants (*M*=0.02, *SD*=0.02; neurotypical>aphasia: *β*=-1.91, *SE*=0.32, *p*<.001) and participants with PD (*M*=0.02, *SD*=0.02; PD>aphasia: *β*=-1.94, *SE*=0.32, *p*<.001). The group by category dimension interactions were not significant for either the neurotypical vs. aphasia comparison (*β*=-.38, *SE*=.21, *p*=.075), nor the PD vs. aphasia comparison (*β*=-0.19, *SE*=0.22, *p*=.381). Follow-up analyses showed no effect of category dimension across groups (*β*=0.06, *SE*=0.25, *p*=.985), nor within the neurotypical group (*β*=0.25, *SE*=0.29, *p*=.732) or the PD group (*β*=0.06, *SE*=0.29, *p*=.990).

#### Response time

RTs in Experiment 2 were the only measure where the pattern was consistent with the LD-specific language recruitment hypothesis. Participants with aphasia were slower to respond during LD trials (*M*=2.37, *SD*=0.70) compared to HD trials (*M*=2.22, *SD*=0.64; LD>HD: *β*=0.16, *SE*=0.08, *p*=.044). The overall RTs for participants with aphasia (*M=*2.30, *SD=* 0.64) were longer than for neurotypical participants (*M=*1.48, *SD=* 0.34; *β*=-.81, SE=0.19, *p*<.001) and participants with PD (*M=*1.43, *SD=*0.29; *β*=-0.86, *SE*=0.19, *p*<.001). We also observed an interaction between group and category dimension for both the neurotypical vs. aphasia comparison (*β*=-0.23, *SE*=0.03, *p*<.001) and the PD vs. aphasia comparison (*β*=-0.19, *SE*=0.03, *p*<.001), such that participants with aphasia had longer RTs for LD categories compared to HD categories. Follow-up analyses showed no overall effect of category dimension across groups (*β*=-0.02, *SE*=0.07, *p*=.985), within the neurotypical group (*β*=0.07, *SE*=0.07, *p*=.669) or within the PD group (*β*=.04, *SE*=0.07, *p*=.907), suggesting that the LD vs. HD difference was specific to participants with aphasia. Note, however, that the effect size of the interactions is much smaller than the overall differences between the aphasia group and the two control groups (**Figure 3C**).

#### Effect of naming performance

To explore the effect of naming ability on categorization performance in this revised version of the task, we again fitted mixed effect regression models with BNT scores, category dimension, interaction between BNT and category dimension, and education level as fixed effects, and participants (across the three groups) and categories as random effects. We found that BNT still significantly predicted all dependent variables (hit rate: *β*=0.38, *SE*=0.13, *p*=.002; false alarm rate: *β*=-0.58, *SE*=0.12, *p*<.001; RT: *β*=-0.29, *SE*=0.07, *p*<.001). As in Experiment 1, this model did not reveal a main effect of category dimension (hit rate: *β*=0.04, *SE*=0.26, *p*=.884; false alarm rate: *β*=-0.12, *SE*=0.26, *p*=.639; RT: *β*=-0.02, *SE*=0.07, *p*=.742); however, unlike Experiment 1, we observed an interaction between BNT and category dimension for hit rate (*β*=0.16, *SE*=0.08, *p*=.039) and RT (*β*=-0.08, *SE*=0.01, *p*<.001). We did not observe such an interaction for false alarm rate (*β*=-0.11, *SE*=0.07, *p*=.090). Finally, education was no longer a significant predictor for any dependent variable (hit rate: *β*=0.11, *SE*=0.15, *p*=.484; false alarm rate: *β*=-0.20, *SE*=0.15, *p*=.190; RT: *β*=-0.01, *SE*=0.08, *p*=.940).

#### Single case analysis

Scrutiny of individual participants’ scores casts some doubt on the relationship between BNT and categorization performance. Specifically, participant A4 in the aphasia group (**Table 4**) had a very low BNT score (1/60), but nonetheless performed very well relative to both the neurotypical and PD groups (hit rate: LD 94%, HD 97%; false alarm rate: LD 2.54%, HD 4.17%; all results are within 2 SD of mean performance in the neurotypical group). This dissociation indicates an absence of a direct causal relationship between naming and categorization.

### 3.3 Interim discussion

Experiment 2 had several goals, including an additional attempt to replicate differences between LD and HD categorization, investigating the effect of severe naming impairments on LD vs. HD categorization performance, testing whether the LD-specific language recruitment hypothesis may find support in a paradigm which is modified to reduce executive demands, and validating a paradigm for use in fMRI. We discuss our results below.

We found no evidence that LD categorization is overall more challenging than HD categorization: no significant differences were observed between LD and HD categories for any of the performance measures (hit rate, false alarm rate, and RT). This result is consistent with Experiment 1 and fails to replicate the results of L&M, who report lower hit rate performance on LD categorization across groups.

As in Experiment 1, naming ability significantly predicted performance on all three outcome measures. Furthermore, possibly because in this experiment we recruited participants with aphasia who had poor naming performance, we also observed a group difference: participants with aphasia had lower hit rates, more false alarms, and longer response times than the two control groups. This evidence points to a possible link between naming performance and categorization. However, as in Experiment 1, this link might arise from the fact that task instructions are presented verbally; thus, linguistic impairments might affect task performance simply because they make it more challenging to process the instructions. Another explanation, also offered by L&M, is that LD categorization is correlated with naming impairments because both tasks may be affected by damage to cognitive control mechanisms, which lay in close proximity to language areas, especially in the left inferior frontal gyrus (LIFG) (e.g., Fedorenko et al., 2012; Kan & Thompson-Schill, 2004; Thompson-Schill et al., 1997). In line with this conjecture, Hu, Small et al. (2021) observed strong neural responses (in fMRI) in the domain-general MD network during an object naming task.

The data from this study offer a further reason to be skeptical about a direct link between naming and categorization. Participant A4 demonstrated an instance of dissociation between these two tasks: despite a BNT score of 1, he performed similar to controls on the categorization task. Dissociations are critical in informing debates about cognitive architecture in general and about the role of language in enabling other cognitive abilities in particular (e.g., Caramazza & Coltheart, 2006). Naturally occurring brain lesions do not respect the boundaries between functionally distinct brain areas, and comorbidities or associations of impairments are common (e.g., E. Bates et al., 2003). For example, damage to the LIFG is likely to cause multiple cognitive impairments due to the high degree of functional heterogeneity of that region (Fedorenko et al., 2012; Fedorenko & Blank, 2020). Thus, as suggested by L&M, a correlation we observe between naming and categorization might be caused by the fact that the brain regions supporting these functions are located nearby (rather than these two functions being supported by the same brain region/mechanism). The dissociation that we observe in participant A4 supports this possibility and suggests that the naming-categorization link might be caused by anatomical coincidence rather than by cognitive interdependency. In his case, severely limited lexical access did not prevent success on the categorization task, revealing that intact linguistic (naming) skills are not necessary for object categorization.

One of the primary goals of Experiment 2 was to establish whether the putative effect of language on LD categorization might manifest more clearly and consistently if the categorization task is modified to reduce the overall cognitive load. We did not find this effect when looking at response accuracies: participants with aphasia did not show a selective impairment in hit rate nor a selective increase in false alarm rate for LD categories. However, we did observe an interaction between RT and category dimension, such that participants with aphasia had longer RTs for LD categories compared to HD categories. We also observed an interaction between naming ability and category dimension, such that lower BNT scores were associated with lower hit rate on LD categories more so than on HD categories, and with longer RTs on LD categories more so than on HD categories. This pattern is consistent with the LD-specific language recruitment hypothesis; however, the fact that we did not observe such a pattern in Experiment 1, which followed L&M’s design and procedure more closely, suggests that this effect is fickle and varies depending on the makeup of the aphasia group and, possibly, the details of the experimental setup. Further, the group by category dimension interaction was absent for the false alarm rate (as previously observed by L&M). Given that, in Experiment 1, the interaction effect for the false alarm rate went in the opposite direction from that predicted by the LD-specific recruitment hypothesis, we find it difficult to reconcile the results from the two experiments with that hypothesis.

The results of Experiments 1 and 2 did not allow us to definitively answer the question of whether language plays a key role in LD categorization. Group comparisons in both experiments failed to replicate the selective LD categorization impairments as reported in L&M; moreover, in Experiment 1, the effect of aphasia on false alarm rate was actually higher for HD categories. On the other hand, Experiment 2 did show a selective decrease in hit rate (and increase in RTs) for LD categories in participants with low naming scores, as predicted by the LD-specific language recruitment hypothesis. However, even this piece of evidence is undermined by the dissociation case of participant A4.

To more definitively establish whether LD categorization recruits the language system, we next turned to fMRI.

## 4. Experiment 3

To further test the relationship between language and categorization, we conducted an fMRI experiment. Neurotypical participants performed the same LD/HD categorization task as participants in Experiment 2. In addition, they completed two ‘localizer’ tasks that were used to identify the networks of interest: the language network and the multiple demand network. The language network responds selectively during language processing, including spoken, written, and signed language comprehension, spoken and written language production, and inner speech (Amit et al., 2017; Braga et al., 2020; Fedorenko et al., 2010, 2011; Giglio et al., 2021; Hu, Small, et al., 2021; Menenti et al., 2011; Scott et al., 2017; Silbert et al., 2014). The multiple demand network is sensitive to general cognitive effort and implicated in cognitive control, responding to a wide range of demanding tasks and exhibiting higher activity when the task is more difficult (Assem, Glasser, et al., 2020; Duncan, 2010; Fedorenko et al., 2013; Hugdahl et al., 2015). Examining activation patterns in both the language and the multiple demand networks allows us to examine the relative contributions of linguistic and cognitive control resources to LD and HD categorization.

As discussed before, brain damage leading to aphasia is often comorbid with multiple demand network damage: the language-selective regions and these domain-general regions in left inferior frontal cortex lie in close proximity to each other (Blank et al., 2014; Fedorenko et al., 2012; Fedorenko & Blank, 2020), with precise locations varying substantially across individuals. Thus, impaired categorization performance of participants with aphasia in Experiments 1 and 2 could have potentially arisen from damage to either network (or to both). Experiment 3 allows us to disambiguate between these possibilities. If, as suggested by L&M, LD categorization indeed relies on language more than HD categorization, we expect to see more activity within the language system during LD trials compared to HD trials. Further, if LD categorization is a more cognitively demanding task, we expect to see higher responses within the multiple demand network during LD trials compared to HD trials (in accordance with the fact that multiple demand regions are sensitive to effort across diverse tasks; Duncan & Owen, 2000; Fedorenko et al., 2013; Hugdahl et al., 2015). Finally, if a brain network does not respond to either LD or HD categorization, we can conclude that this network is not recruited for this task.

### 4.1 Method

#### 4.1.1 Participants

Fourteen neurotypical participants (7 F, age *M*=22.31, *SD*=3.51) were recruited from MIT and the surrounding community and paid $60 for their participation. All were native speakers of English. One participant was left-handed (see Willems et al., 2014, for motivation to include left-handers in cognitive neuroscience research) but showed typical left-lateralized language activation as determined by the language localizer task (described below). All participants gave informed consent in accordance with the requirements of MIT’s Committee On the Use of Humans as Experimental Subjects (COUHES).

#### 4.1.2 Design, materials, and procedure

Each participant completed a language localizer task aimed at identifying language-responsive brain regions (Fedorenko et al., 2010), a spatial working memory (WM) task aimed at identifying the multiple demand network (Fedorenko et al., 2013), and the critical categorization task. Some participants completed one or more additional tasks for unrelated studies. The entire scanning session lasted two hours.

##### Language network localizer

Participants read sentences (e.g., NOBODY COULD HAVE PREDICTED THE EARTHQUAKE IN THIS PART OF THE COUNTRY) and lists of unconnected, pronounceable nonwords (e.g., U BIZBY ACWORRILY MIDARAL MAPE LAS POME U TRINT WEPS WIBRON PUZ) in a blocked design. Each stimulus consisted of twelve words/nonwords. The sentences > nonword-lists contrast has been previously shown to reliably activate high-level language processing regions and to be robust to changes in the materials, task, and modality of presentation (Fedorenko et al., 2010; Mahowald & Fedorenko, 2016; Scott et al., 2017). For details of how the language materials were constructed, see Fedorenko et al. (2010). The materials are available at http://evlab.mit.edu/funcloc. Stimuli were presented in the center of the screen, one word/nonword at a time, at the rate of 450ms per word/nonword. Each stimulus was preceded by a 100ms blank screen and followed by a 400ms screen showing a picture of a finger pressing a button, and a blank screen for another 100ms, for a total trial duration of 6s. Participants were asked to press a button whenever they saw the picture of a finger pressing a button. This task was included to help participants stay alert and awake. Condition order was counterbalanced across runs. Experimental blocks lasted 18s (with 3 trials per block), and fixation blocks lasted 14s. Each run (consisting of 5 fixation blocks and 16 experimental blocks) lasted 358s. Each participant completed 2 runs.

##### Multiple demand network localizer

Participants had to keep track of four (easy condition) or eight (hard condition) sequentially presented locations in a 3×4 grid (Fedorenko et al., 2013). The hard > easy contrast has been previously shown to robustly activate multiple demand regions (Assem, Blank, et al., 2020; Blank et al., 2014; Fedorenko et al., 2013; Mineroff et al., 2018). Stimuli in both conditions were presented in the center of the screen across four steps. Each of these steps lasted for 1s and presented one location on the grid in the easy condition, and two locations in the hard condition. Each stimulus was followed by a choice-selection step, which showed two grids side by side. One grid contained the locations shown on the previous four steps, while the other contained an incorrect set of locations. Participants were asked to press one of two buttons to choose the grid that showed the correct locations. Condition order was counterbalanced across runs and participants. Experimental blocks lasted 32s (with 4 trials per block), and fixation blocks lasted 16s. Each run lasted 448s, consisting of 12 experimental blocks (6 per condition), and 4 fixation blocks. Twelve participants completed two runs and two participants completed one run.

##### Critical categorization task

The categorization materials were the same as those used in Experiment 2 (see **Figure 1, bottom**). The timing differed in the following way. In order to make blocks uniform in duration, each category block started with a category label presented for 2s, and then the 12 images were presented sequentially at the fixed speed of 2s per image. As in Experiment 2, any given category block contained between 4 and 6 target images. Participants were asked to press a button if the picture belonged to the target category and not to press anything if it did not. As before, the category label was displayed at the top of the screen for the duration of the trial to minimize memory demands. Category blocks lasted 26s (2s category label presentation + 2s * 12 images), and fixation blocks lasted 14s. Each run, consisting of 12 category blocks (6 LD and 6 HD) and 4 fixation blocks, lasted 368s. Each participant completed 3 runs. Across the 3 runs, any given participant saw a random subset of the 32 categories, with some categories repeating (but never repeating within a run; see **Appendix 1 Table 1** for details). Condition order was counterbalanced across runs and participants.

#### 4.1.3 fMRI data acquisition

Structural and functional data were collected on the whole-body, 3 Tesla, Siemens Trio scanner with a 32-channel head coil, at the Athinoula A. Martinos Imaging Center at the McGovern Institute for Brain Research at MIT. T1-weighted structural images were collected in 176 sagittal slices with 1mm isotropic voxels (TR=2530ms, TE=3.48ms). Functional, blood oxygenation level dependent (BOLD), data were acquired using an EPI sequence (with a 90° flip angle and using GRAPPA with an acceleration factor of 2), with the following acquisition parameters: thirty-one 4mm thick near-axial slices acquired in the interleaved order (with 10% distance factor), 2.1mm×2.1mm in-plane resolution, FoV in the phase encoding (A>>P) direction 200mm and matrix size 96mm×96mm, TR=2000ms and TE=30ms. The first 10s of each run were excluded to allow for steady state magnetization.

#### 4.1.4 fMRI data preprocessing

MRI data were analyzed using SPM12 and custom MATLAB scripts (available in the form of an SPM toolbox from https://evlab.mit.edu/funcloc/). Each participant’s data were motion corrected (realignment to the mean image of the first run using 2nd-degree b-spline interpolation) and then normalized into a common brain space (the Montreal Neurological Institute (MNI) template) (estimated for the mean image using trilinear interpolation) and resampled into 2mm isotropic voxels. The data were then smoothed with a 4mm FWHM Gaussian filter and high-pass filtered (at 128s). Effects were estimated using a General Linear Model (GLM) in which each experimental condition was modeled with a boxcar function (modeling entire blocks) convolved with the canonical hemodynamic response function (HRF). The model also included first-order temporal derivatives of these effects, as well as nuisance regressors representing entire experimental runs, offline-estimated motion parameters, and timepoints classified as outliers based on the motion parameters.

#### 4.1.5 Defining individual functional regions of interest (fROIs)

Responses to the critical categorization experiment were extracted from regions of interest that were defined functionally in each individual participant (Nieto-Castañón & Fedorenko, 2012; Saxe et al., 2006). Three sets of functional regions of interest (fROIs) were defined—one for the language network, one for the multiple demand network, and one for the putative LD>HD categorization regions. To do so, we used the Group-constrained Subject-Specific (GSS) approach (Fedorenko et al., 2010; Julian et al., 2012). In particular, fROIs were constrained to fall within a set of “parcels”, which marked the expected gross locations of activations for the relevant contrast. For the language network, the parcels were generated based on a group-level representation of language localizer data from 220 participants. For the multiple demand network, the parcels were generated based on a group-level representation of spatial working memory task data from 197 participants. For the putative LD categorization regions, we generated the parcels based on the data collected in this study. The parcels are available on OSF (https://osf.io/guwh8/). To create each set of parcels, individual activation maps for the relevant localizer contrast were binarized (by turning all voxels significant at the *p*<.001 whole-brain threshold (uncorrected) into 1s, and the rest into 0s) and overlaid in the MNI space to create a probabilistic overlap map. Note that for the multiple demand network, the individual activation maps were averaged across the two hemispheres prior to binarizing. The map was then smoothed (FWHM = 6mm), and voxels with fewer than 10% of participants overlapping were excluded. The resulting map was divided into regions using a watershed algorithm. Finally, we excluded parcels that did not show significant effects for the relevant localizer contrast in a left-out run or did not contain supra-threshold voxels in at least 60% of the participants (for language and multiple demand networks) or in at least 50% of the participants (for putative LD categorization regions). For the multiple demand network, we also a) excluded parcels in the visual cortex (the hard condition includes more visual information than the easy condition and thus yields more activation in the visual cortex), and b) divided a parcel that encompassed parts of both the precentral gyrus and the opercular portion of the inferior frontal gyrus according to the macroanatomical boundary.

For each participant, each set of masks was intersected with the participant’s activation map for the relevant contrast (sentences>nonwords for the language network, hard>easy spatial WM for the multiple demand network, and LD>HD for putative LD categorization regions). Within each mask, the voxels were sorted based on their *t* values for the relevant contrast, and the top 10% of voxels were selected as that participant’s fROI. This top n% approach ensures that the fROIs can be defined in every participant, thus enabling us to generalize the results to the entire population (Nieto-Castañón & Fedorenko, 2012).

#### 4.1.6 Examining the functional response profiles of fROIs

After defining fROIs in individual participants, we evaluated their responses to the conditions of interest by averaging the responses across voxels to get a single value per condition per fROI. The responses to the localizer conditions (sentences and nonwords for language fROIs, hard and easy working memory conditions for multiple demand fROIs, and LD and HD categorization for categorization fROIs) were estimated using an across-runs cross-validation procedure, where one run was used to define the fROI and the other to estimate the response magnitudes, then the procedure was repeated switching which run was used for fROI definition vs. response estimation, and finally the estimates were averaged to derive a single value per condition per fROI per participant. This cross-validation procedure allows one to use all of the data for defining the fROIs as well as for estimating their responses (see Nieto-Castañón & Fedorenko, 2012, for discussion), while ensuring the independence of the data used for fROI definition and response estimation (Kriegeskorte et al., 2009). Two participants completed only one run of the multiple demand localizer task; therefore, we did not estimate the strength of their responses to the hard and easy multiple demand localizer conditions but ensured that the whole-brain activation maps for the hard>easy contrast looked as expected.

#### 4.1.7 Statistical analyses

Similar to Experiments 1 and 2, we analyzed our data using mixed effect regression models (Baayen et al., 2008). For hit rate and false alarms we use logistic regression (Jaeger, 2008). For RT and fROIs response magnitudes, we use linear regression. In all models, condition was a fixed effect and participant was a random intercept. The model for the multiple demand network included hemisphere as an additional fixed effect. For language and multiple demand network analyses, we also included fROI as a random intercept and then ran follow-up analyses on individual fROIs using false discovery rate (FDR) correction (Benjamini & Hochberg, 1995) for the number of fROIs in each network. Behavioral analyses used sum coding for condition (LD vs. HD in the categorization task and Hard vs. Easy in the multiple demand localizer task). Neuroimaging analyses used custom contrasts (see **Appendix 2** for detailed contrast specification). The mixed effect analyses were run using the *lmer* function from the *lme4* R package (D. Bates et al., 2015); statistical significance of the effects was evaluated using the *lmerTest* package (Kuznetsova et al., 2017). The hypotheses-specific contrasts were defined using the *hypr* package (Rabe et al., 2020).

If linguistic resources are engaged during categorization, we would expect an overall high response of the language network to categorization conditions. Further, if, as L&M have argued, LD categorization taxes linguistic resources to a greater extent, we would expect to see stronger response of this network to the LD compared to the HD condition. Lastly, if LD categorization is generally more taxing, we would expect to see greater responses to the LD condition in the domain-general multiple demand regions that are sensitive to effort across diverse tasks (Duncan, 2010, 2013; Fedorenko et al., 2013; Hugdahl et al., 2015).

### 4.2 Results

#### 4.2.1 Behavioral data

##### Multiple demand network localizer

Due to a technical error, behavioral data for one participant got overwritten. For the remaining thirteen participants, performance on the spatial working memory task was as expected: participants were more accurate and faster in the easy condition (accuracy *M*=93.91%, *SD*=3.00%; reaction time (RT)=1.18s, *SD*=0.16s) than the hard condition (accuracy *M*=79.65%, *SD*=12.03%; RT *M*=1.52s, *SD*=0.25s). Mixed effect models with condition as a fixed effect and participant as a random intercept showed that both accuracy and RT effects were significant (accuracy: *β* = -1.41, SE=0.202, *p* < .001; RT: *β* =0.33, *SE* =0.027, *p* < .001).

##### Critical categorization task

The hit rate for the two categorization conditions was not significantly different (LD *M* =93.62%, *SD* =9.84%; HD *M* =93.42%, *SD* =8.65%; LD>HD *β* =0.03, *SE* =0.22, *p* =.897). Similarly, the number of false alarms did not significantly differ across conditions (LD *M* =3.01%, *SD* =1.43%; HD *M* =3.34%, *SD* =2.39%; LD>HD *β* = -0.37, *SE* =0.38, *p* =.329). Finally, there was no significant difference between response times in the LD condition (RT=0.81s, *SD* =0.1s) and the HD condition (RT=0.84s, *SD* =0.1s; LD>HD *β* = -0.03, *SE* =0.02, *p* =.156).

#### 4.2.2 Functional response profile of the language network

Although the sentence reading condition elicited strong responses in the language fROI, the responses to the categorization task were only marginally above 0 (*β* =0.42, *SE* =0.19, *p* =.054; see **Figure 4**), not significantly different from responses to nonword reading (*β* =0.13, *SE* =0.09, *p* =.144), and significantly weaker than responses to sentences (*β* = -1.49, *SE* =.09, *p*<.001). Further, there was no significant difference between responses to LD and HD categorization (*β* = -0.02, *SE* =0.10, *p* =.848).

**Figure 4.**
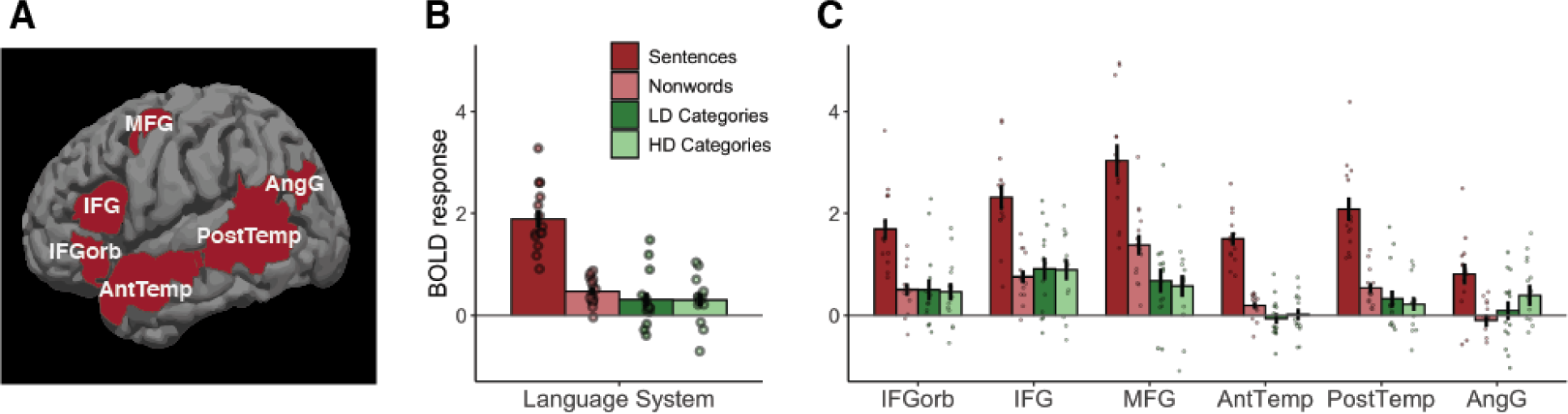
Categorization responses within the language brain network. (**A**) Parcels used to define functional regions of interest (fROIs) in individual participants. (**B**) Average responses within the language network to four conditions of interest (sentence reading and nonword reading vs. LD and HD categorization). (**C**) fROI responses to the four conditions of interest.

Follow-up analyses in individual language fROIs (**Appendix 2 Table 1**) showed that responses to categorization were significantly above 0 in frontal fROIs (MFG, IFG, and IFGorb). However, none of the responses were significantly higher than responses during the control task, nonword reading, indicating that these responses are not language-specific. Thus, our results suggest that the language network is not involved in either LD or HD categorization in neurotypical participants.

#### 4.2.3 Functional response profile of the multiple demand network

Multiple demand network responses to categorization were significantly above 0 (*β* =1.07, *SE* =0.21, *p*<.001; see **Figure 5**) and stronger than responses to control conditions from the language localizer task (categorization > sentences: *β* =0.73, *SE* =0.08, *p*<.001; categorization > nonwords: *β* =0.41, *SE* =0.08, *p*<.001). However, they were weaker than responses to the spatial working memory task (*β* = -1.43, *SE* =.07, *p*<.001), indicating that the working memory task was more effortful. The responses to the categorization task were stronger in the left hemisphere (*β* =0.24, *SE* =0.09, *p* =.005). We also observed an interaction between the working memory > categorization contrast and hemisphere (*β* =0.29, *SE* =0.13, *p* =.024), showing that the working memory task engages the right hemisphere to a greater extent. There was also an interaction between the Hard>Easy working memory task and hemisphere, such that the effect was greater in right hemisphere (*β* =0.38, *SE* =0.19, *p* =.040).

**Figure 5.**
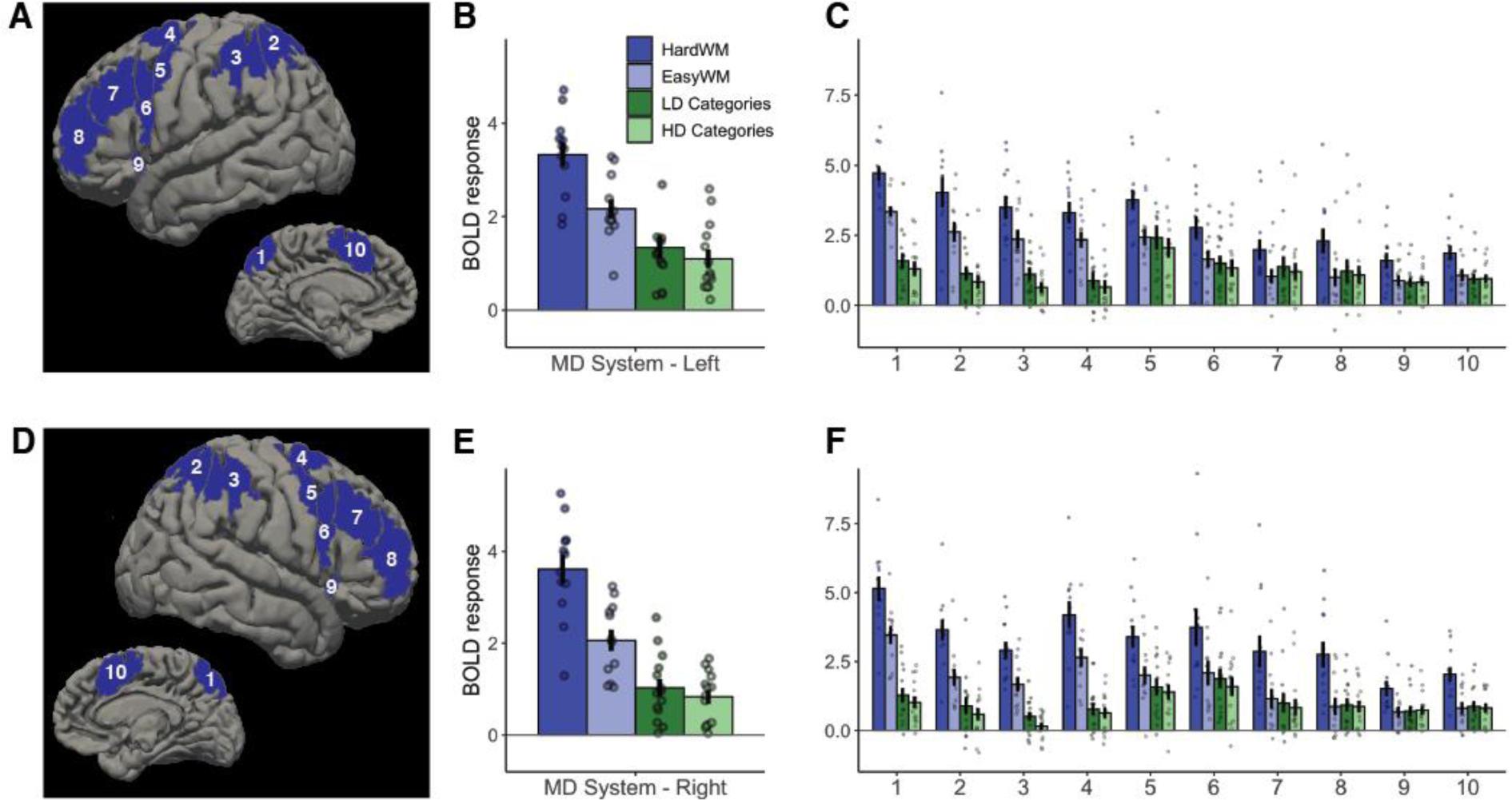
Categorization responses within the multiple demand brain network. (**A**) Left hemisphere parcels used to define functional regions of interest (fROIs) in individual participants. (**B**) Average responses within the left hemisphere fROIs to four conditions of interest (hard and easy working memory tasks vs. LD and HD categorization). (**C**) Left hemisphere fROI responses to the four conditions of interest. (**D-F**) Parcels, average responses, and fROI-level responses in the right hemisphere.

Importantly, there was a small but significant difference between responses to LD and HD categorization tasks (*β* =0.19, *SE* =0.09, *p* =.025), indicating that LD categorization is slightly more effortful than HD categorization in neurotypical participants.

Follow-up analyses on individual fROIs (**Appendix 2 Table 2**) showed that responses to categorization were significantly above 0 in all fROIs. However, they were weaker than the overall responses to the working memory task in almost all fROIs (except left middle frontal fROI). This result highlights the domain-general nature of these responses. Further, none of the fROIs had significantly different responses to LD and HD categories, despite the presence of this effect in the network-level analysis.

#### 4.2.4 Whole-brain analyses

We also conducted a whole-brain analysis to identify fROIs that might respond more strongly to LD or HD categorization but lie outside the language and multiple demand fROIs described above. The GSS analysis (see **Methods** for details) revealed that no regions exhibited consistent HD>LD responses across participants; however, the LD>HD contrast revealed two parcels, both located in left parietal lobe (**Figure 6**). Further analysis of fROIs defined within these parcels showed that the LD>HD response only reached significance in fROI 2 (*β* =0.43, *SE* =0.17, *p* =.013), but not in fROI 1 (*β* =0.58, *SE* =0.30, *p* =.060). The overall categorization response was significantly above 0 in fROI 1 (*β* =0.65, *SE* =0.19, *p* =.001) but not fROI 2 (*β* = -0.13, *SE* =0.15, *p* =.389).

**Figure 6.**
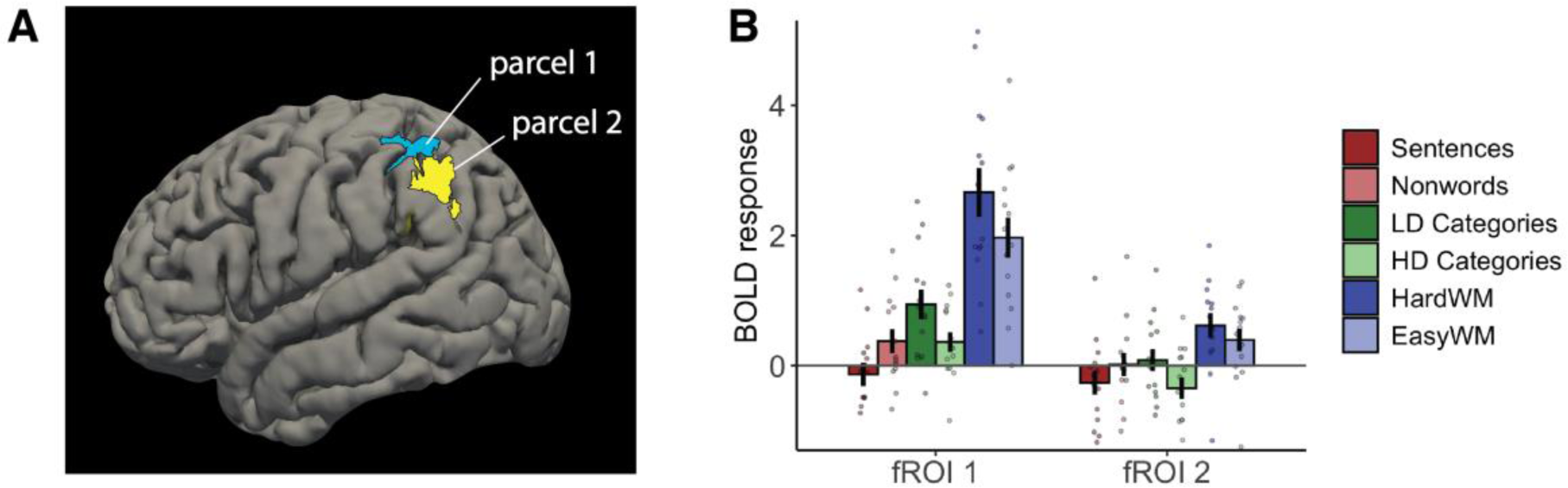
Results of the whole-brain analyses. (A) Parcels defined with the LD>HD categorization contrast. (B) Responses to conditions of interest within the two fROIs (defined as the top 10% of voxels within each parcel, sorted by the magnitude of the LD>HD response). WM – working memory task.

Importantly, both fROIs responded to the working memory task more strongly than to the categorization task (fROI 1: *β* =1.66, *SE* =0.21, *p*<.001; fROI 2: *β* =0.64, *SE* =0.12, *p*<.001), indicating that these regions likely respond to general cognitive effort rather than to LD categorization (or feature selection) specifically, and thus likely belong to the MD network. Neither of the two fROIs exhibited a sentences>nonwords effect (fROI 1: *β* = -0.51, *SE* =0.30, *p* =.094; fROI 2: *β* = -0.28, *SE* =0.17, *p* =.098), which shows that these regions do not respond to linguistic input.

The whole-brain analysis provides additional evidence against the LD-specific language recruitment hypothesis and shows that differences in LD vs. HD categorization, if present, are likely caused by domain-general mechanisms.

### 4.3 Interim discussion

In Experiment 3, we used fMRI to examine neural responses to LD and HD categorization. Our main goal was to evaluate the hypothesis that LD categorization relies more heavily on linguistic resources compared to HD categorization. For this purpose, we identified the language network individually in 14 healthy adults and examined its responses during LD and HD categorization. The language network exhibited low responses to both categorization tasks, which did not differ from activations elicited by reading of nonword sequences (a low-level control condition). There was no difference between responses to LD categories and responses to HD categories, contra the prediction that the language network would be selectively or preferentially engaged during LD categorization.

Unlike the language network, the domain-general multiple demand network (also defined individually in each participant) was engaged during categorization, indicating that this task is cognitively challenging. This network responded more strongly to LD than HD categorization, but this effect was small. The whole-brain analyses specifically aimed at identifying regions with stronger responses to LD than HD categorization confirmed these results: both fROIs it identified responded more strongly to a working memory task than to categorization task, and the LD>HD effect was small and/or not statistically significant. We conclude that categorization, and LD categorization in particular, relies on domain-general multiple demand regions and not on language-specific regions.

Neuroimaging of healthy individuals provides a powerful complement to patient studies. Given the strong and selective engagement of the language network during all behaviors requiring access to linguistic representations (Fedorenko et al., 2010, 2011; Giglio et al., 2021; Hu et al., 2021; Menenti et al., 2011; Scott et al., 2017, among others), the lack of activity in the language regions during categorization strongly suggests that they do not contribute to categorization (Mather et al., 2013). The response to categorization within the multiple demand network, on the other hand, indicates its involvement in categorization, even though we note that fMRI evidence described here is correlational, not causal, and should be complemented with patient studies or brain stimulation studies that specifically target this hypothesis. Examining neuroimaging evidence is particularly helpful when patient studies do not produce conclusive results, as in our case.

Whereas some previous work suggested that a region within left angular gyrus is involved in inhibiting irrelevant semantic information (Lewis et al., 2019), as may be required for LD categorization, the results of our study suggest that activation of the language-responsive portion of the left angular gyrus was comparable during LD and HD categorization. If anything, this language fROI showed numerically higher activation during HD categorization, suggesting that it may be recruited for recognizing and thinking about established sets more than for constructing novel sets that may require inhibition of object-irrelevant characteristics. We also did not find significant differences in the engagement of the language fROIs in the left inferior frontal cortex during LD and HD categorization. These results are in contrast to findings from Lupyan et al. (2012), which suggested that tDCS to the left inferior frontal cortex disrupted performance on LD but not HD categorization. This may be because the left inferior frontal cortex contains not only language-responsive areas, but also multiple demand areas (Fedorenko et al., 2012; Fedorenko & Blank, 2020), and interfering with the latter areas’ activity may have a disproportionately higher effect on LD categorization.

The response to categorization within the multiple demand network was stronger in the left hemisphere, consistent with the view that label-based categorization recruits the left hemisphere more strongly (e.g., Franklin et al., 2008; Gilbert et al., 2006). This makes the categorization task similar to logic and math, which also evoke left-lateralized responses within the multiple demand network (Amalric & Dehaene, 2016; Monti et al., 2009, 2012; Pinel & Dehaene, 2009). Importantly, our result demonstrates that, just because the function is left-lateralized, it is not necessarily related to language, at least not in fully formed brains (contra, e.g., Gilbert et al., 2006; see also Holmes & Wolff, 2012).

All in all, results from Experiment 3 disconfirm the hypothesis that LD categorization relies on linguistic resources. Instead, they show that categorization recruits the multiple demand brain regions and that LD categorization is, on average, slightly more effortful that HD categorization.

## 5. Discussion

We reported three experiments that evaluated the hypothesis that linguistic resources are essential for performing feature-based, or low-dimensional (LD), categorization—what we refer to as the ‘LD-specific language recruitment hypothesis’ (Langland-Hassan et al., 2021; Lupyan, 2009; Lupyan et al., 2012; Lupyan & Mirman, 2013). In Experiment 1, we aimed to replicate the results of Lupyan and Mirman (2013), who showed a selective impairment in LD categorization in individuals with aphasia. Our results failed to replicate this critical finding, although they did show that naming ability, as measured by Boston Naming Test (BNT) scores, was a significant predictor of overall categorization performance.

In Experiment 2, we modified the design to reduce general task complexity and examined the specific contribution of naming ability to categorization by recruiting a group of participants with very low naming scores. Similar to Experiment 1, we found no significant interaction between participant group and LD/HD categorization. However, we did find a significant interaction between BNT and LD/HD categorization for one of the two accuracy measures (hit rate). Although this result lends some support to the LD-specific language recruitment hypothesis, a case-by-case analysis identified an individual with a severe naming impairment (with a score of 1 out of 60 on the BNT) who performed within the neurotypical range on both HD and LD categorization. Evidence from patients with brain lesions remains an important way to establish whether specific cognitive capacities support performance on particular tasks (Rorden & Karnath, 2004). Such studies have previously demonstrated that many high-order cognitive functions are not affected even in the presence of severe linguistic deficits (e.g., Apperly et al., 2006; Bek et al., 2013; Chen, Affourtit et al., 2021; Varley et al., 2001, 2005; Varley & Siegal, 2000; Willems et al., 2011). Based on Experiment 2, we therefore concluded that lexical retrieval is *not necessary* for successful categorization, including categorization based on single features.

In Experiment 3, we used a complementary approach and examined the engagement of the language network and a domain-general multiple demand network in HD and LD categorization using fMRI in neurotypical adults. The language network was not engaged during either LD or HD categorization: its responses did not significantly differ from responses during the control, nonword reading, task. This observation goes against the hypothesis that categorization (either LD or HD) relies on linguistic resources. In contrast, the multiple demand network was recruited during the categorization task, consistent with prior evidence of its involvement in diverse cognitively challenging tasks (Assem, Glasser, et al., 2020; Duncan, 2010, 2013; Fedorenko et al., 2013). It also responded more strongly during LD than HD categorization. Given extensive evidence that the multiple demand network responds more strongly when the task is harder (e.g., Fedorenko et al., 2011, 2013; Hugdahl et al., 2015; Shashidhara et al., 2019), the increased response during LD categorization is consistent with the hypothesis that LD categorization is more cognitively challenging. However, this effect was small and did not come out as statistically significant in any of the individual multiple demand regions in follow-up analyses. In sum, we find little evidence in favor of the LD-specific language recruitment hypothesis.

### 5.1 The cognitive control account of categorization performance

The failure to replicate the results from L&M in Experiment 1 and an only partial replication in Experiment 2 have several possible explanations. The first explanation is that the effect described by L&M is real, but we could not detect it due to low power (e.g., small sample size). This explanation is unlikely because of our neuroimaging results: if language was indeed required for LD categorization, the language network would be active during the LD categorization condition. The second explanation is that the result that was reported by L&M is a false positive. The third explanation is that the effect holds in a subset of individuals with aphasia, due to comorbid cognitive control impairments. We cannot definitively rule out either the second or the third explanation, although our neuroimaging results provide some support for the latter: the multiple demand network, implicated in cognitively demanding tasks, was somewhat more active during LD than during HD categorization.

The hypothesis that cognitive control deficits underlie impaired categorization can also explain the link between categorization and naming, which we observed in both Experiments 1 and 2, and which was also reported by L&M. Confrontation naming is a complex, multi-component behavior that involves not only linguistic, but also visual, motor-articulatory, and critically, executive resources. Indeed, a recent fMRI study (Hu, Small, et al., 2021) reports strong responses within the multiple demand network to an object naming condition. Furthermore, unlike syntactic comprehension, both naming ability and fluid intelligence (a trait linked to the multiple demand network; Gläscher et al., 2010; Woolgar et al., 2010, 2018) decline with age, and this decline is linked to decreased activity in the multiple demand brain regions during both of these tasks (Samu et al., 2017). Thus, although both our work and L&M show the relationship between naming and categorization, the underlying cognitive mechanism of this relationship is likely related to cognitive control, not language.

Future patient studies should explicitly test the cognitive control account of LD-selective categorization impairments. One way to do so is to use lesion mapping along with probabilistic maps of language and multiple demand networks (see, e.g., Woolgar et al., 2018): this method allows explicitly determining which of these two networks underlies observed behavior patterns. Another way is to measure cognitive control in individuals with brain damage and use it as a predictor when evaluating the relationship between naming performance and categorization. Yet another approach would be to explore these relationships in neurotypical individuals by examining the correlational structure of these abilities. Such studies could provide additional evidence in favor or against the cognitive control account of categorization impairments, complementing our neuroimaging results and reconciling conflicting findings from individuals with aphasia.

### 5.2 The relevance of LD vs. HD distinction

Why did we find no, few, or inconsistent differences in performance and neural responses between LD and HD categories? A possible explanation is that “LD” and “HD” category types are not ‘natural kinds’. As discussed in the introduction, different researchers have emphasized different distinctions among categories, such as natural/ad hoc, taxonomic/thematic, dense/sparse, concrete/abstract, etc. Many of these distinctions are not isomorphic with the LD/HD distinction. In particular, HD categories encompass both taxonomic (e.g., “animals”) and thematic (e.g., “non-food things found in the kitchen”) categories. Multiple studies show that the processing of taxonomic and thematic relations relies on distinct cognitive and neural mechanisms (e.g., Kalénine et al., 2009; Lewis et al., 2015; Sass et al., 2009; Schwartz et al., 2011; Xu et al., 2018); collapsing them into a single “HD” category type leads to substantial within-HD heterogeneity and may therefore obscure potential HD/LD differences.

Furthermore, not all LD categories as defined by Lupyan and Mirman (2013) necessarily involve conceptual processing. For instance, many are based on color: e.g., “THINGS THAT ARE YELLOW”. Although color is often encoded as part of the conceptual representation of an object, this conceptual representation was not required for the task in question: participants were simply asked to indicate whether the object they were viewing was yellow, and decisions could be made on the basis of surface perceptual features alone. Thus, even if ‘true’ (semantic) LD categories are indeed harder to process than HD categories, inclusion of perception-based color categories could have prevented us from reliably observing this difference.

Our results are somewhat inconsistent with recent work by Langland-Hassan et al. (2021), who observe that individuals with aphasia were slower when processing abstract categories compared to concrete categories. The authors argue that the abstract/concrete distinction is similar to the LD/HD distinction because members of abstract categories share fewer common features. However, another important difference is the *kind* of features used for categorization. For instance, their example of an abstract category “predict” (which includes a weatherperson and a fortune-teller) relies on an unobservable functional similarity rather than on an observable visual similarity. Unobserved features play an important role in the use of verbal category labels (Gelman & Roberts, 2017), so it is possible that language mediates categorization based on *latent* features rather than LD categorization per se. In short, the LD/HD and the abstract/concrete distinction do not cleanly map onto each other, which makes it difficult to compare the results of our experiments to those by Langland-Hassan et al. More generally, the typology of category types remains vague and inconsistent, and more careful work should be done to establish meaningful category distinctions and thus facilitate comparisons across studies.

### 5.3 Possible paradigm-specific effects of verbal labels

Even if we were able to successfully replicate L&M’s findings, our conclusions about the language–categorization link would be complicated by the fact that the paradigm introduced by L&M is not language-free. In order to successfully sort objects into categories, participants need to read (or hear) and encode the category label, presented verbally. The importance of language during the instruction encoding stage might account for the relationship between categorization performance and naming ability; it might even explain the (putative) LD-specific categorization impairments, given that category labels for LD categories are often longer. In Experiments 2 and 3, we modified the experimental paradigm to temporally separate the instruction processing step and the categorization step, which allowed us to measure the behavioral and neural responses to categorization alone. Another solution to this issue would be to modify the paradigm to remove verbal labels altogether, e.g., by providing several category exemplars instead.

In addition, linguistic labels might contribute to the task via verbal rehearsal: participants might employ a phonological loop to maintain an active representation of the labels in working memory. Such assistive role of language labels has been observed in conditions of high cognitive demand (e.g., during mathematical calculation; Benn et al., 2012; Klessinger et al., 2012). However, such low-level verbal/phonological rehearsal appears to rely on lower-level speech processing mechanisms (e.g., Scott & Perrachione, 2019) and the domain-general multiple-demand network (e.g., Fedorenko et al., 2011; Shashidhara et al., 2020), not on the language network. In any case, the verbal rehearsal account is quite different from L&M’s original LD-specific language recruitment hypothesis.

### 5.4 Relationship to other work on language and categorization

Other results from psycho- and neurolinguistics also support the view that linguistic resources do not typically mediate categorization in humans. If access to linguistic representations were necessary for categorization, categorizing images would take longer than categorizing words; instead, they take approximately the same amount of time (Potter & Faulconer, 1975). When asked to match a picture with a label, participants do not explicitly generate/rehearse verbal labels in advance unless there is an additional memory demand (e.g., if images disappear from the screen) (Pontillo et al., 2015). Previous work also shows that language is not necessary for performing tasks that require isolating a specific aspect (“feature”) of the semantic representation, including theory of mind inferences (Varley et al., 2001; Varley & Siegal, 2000) and thematic role identification (Ivanova et al., 2021). Our work therefore adds to the growing body of evidence for a separation between linguistic and visual semantic processing.

That said, many studies have shown that *linguistic labels* influence categorization behavior in infants (e.g., Ferguson & Waxman, 2017; Gershkoff-Stowe et al., 1997; Plunkett et al., 2008; Sloutsky & Fisher, 2004; Waxman & Gelman, 2009) and adults (e.g., Brojde et al., 2011; Lupyan, 2009; Lupyan et al., 2007; Zettersten & Lupyan, 2020), so the relationship between words and categories is clearly an important one. What we are showing here is that the mechanisms responsible for language processing are not engaged during object categorization, nor are they specifically recruited for LD categorization. It is possible that linguistic labels, once acquired, may influence categorization via other brain systems, e.g., semantic, domain-general, or perceptual. The cognitive and neural mechanisms underlying the influence of labels on categorization thus remain to be determined (see, e.g., Gliozzi et al., 2009; Ivanova & Hofer, 2020; Luo et al., 2021; Lupyan, 2012, for some modeling proposals).

Overall, our study shows that categorizing items is not a language-dependent task in the adult brain, regardless of whether the categorization is made on the basis of multiple features (HD) or a single feature (LD). Instead, this task relies on the domain-general multiple demand system, which supports diverse goal-directed behaviors. Our work provides evidence against the view of language as an aid for feature-based (LD) categorization and highlights the value of complementing patient studies with neuroimaging experiments.

## Footnotes

1 In some places, Lupyan and colleagues talk about ‘forming the task-relevant category representations’ (e.g., Lupyan & Mirman, 2013, p. 1188). However, this construal is confusing given that in the paradigms used in these studies, the category label is always provided to the participants. As a result, we assume that the argument has to revolve around the *maintenance* of task-relevant information.

2 We will adhere to the ‘LD’/‘HD’ terminology in the remainder of the paper for consistency with Lupyan and colleagues’ work.

3 L&M state that they only included ‘the correct responses’ in their RT analyses. It is not clear what is meant here given the internal complexity of the trials (i.e., possible errors including misses and false alarms). It is possible that L&M only included trials where *no errors of any kind* were made, but they also talk about ‘per click’ RTs, which is not consistent with this interpretation. It also appears that L&M analyzed median, not mean RTs. For simplicity and to avoid collider bias (Elwert & Winship, 2014), we chose to analyze all trials here. We use mean per-trial values, but we make the per-image data available on OSF (https://osf.io/guwh8/), so other researchers could perform additional analyses.

