## Appendix 1 for "No evidence for a special role of language in feature-based categorization"

**Appendix 1: Experiment 3, Material Presentation Details**

**Table 1.** Distribution of category use across participants (i.e., the number of times each participant saw each category during the categorization experiment, summed across runs).

|  |  | **Subject ID** | | | | | | | | | | | | | |
| --- | --- | --- | --- | --- | --- | --- | --- | --- | --- | --- | --- | --- | --- | --- | --- |
| **Condition** | **Category** | **1** | **2** | **3** | **4** | **5** | **6** | **7** | **8** | **9** | **10** | **11** | **12** | **13** | **14** |
| HD | animals that live in water | 1 | 1 | 1 | 0 | 0 | 0 | 1 | 1 | 2 | 1 | 1 | 2 | 2 | 2 |
| HD | birds | 2 | 0 | 1 | 1 | 1 | 3 | 1 | 2 | 1 | 2 | 3 | 1 | 3 | 1 |
| HD | clothes | 1 | 1 | 1 | 1 | 1 | 0 | 1 | 0 | 0 | 2 | 1 | 0 | 1 | 1 |
| HD | dangerous animals | 2 | 1 | 2 | 1 | 1 | 0 | 1 | 1 | 1 | 1 | 1 | 0 | 0 | 1 |
| HD | farm animals | 1 | 1 | 3 | 2 | 0 | 0 | 2 | 2 | 2 | 1 | 3 | 1 | 2 | 1 |
| HD | fruit | 1 | 2 | 1 | 1 | 2 | 2 | 0 | 0 | 1 | 1 | 1 | 2 | 1 | 0 |
| HD | home appliances | 2 | 2 | 0 | 1 | 1 | 2 | 2 | 1 | 0 | 1 | 0 | 2 | 0 | 1 |
| HD | insects | 2 | 2 | 1 | 3 | 1 | 2 | 1 | 2 | 1 | 2 | 1 | 2 | 1 | 2 |
| HD | musical instruments | 0 | 1 | 1 | 0 | 3 | 0 | 2 | 1 | 2 | 2 | 2 | 1 | 1 | 1 |
| HD | non food things found in the kitchen | 2 | 1 | 0 | 0 | 2 | 2 | 0 | 2 | 2 | 1 | 1 | 2 | 0 | 1 |
| HD | objects found in the laundry room | 1 | 1 | 0 | 2 | 1 | 1 | 1 | 2 | 1 | 1 | 1 | 0 | 1 | 1 |
| HD | objects that hold water | 0 | 1 | 2 | 2 | 0 | 1 | 2 | 0 | 0 | 1 | 1 | 1 | 2 | 2 |
| HD | objects used for transportation | 1 | 2 | 1 | 1 | 1 | 1 | 0 | 1 | 3 | 1 | 0 | 0 | 1 | 1 |
| HD | things that fly | 1 | 2 | 3 | 1 | 0 | 1 | 2 | 2 | 1 | 0 | 1 | 2 | 0 | 2 |
| HD | tools | 1 | 0 | 0 | 2 | 2 | 2 | 2 | 0 | 0 | 0 | 0 | 1 | 0 | 0 |
| HD | vegetables | 0 | 0 | 1 | 0 | 2 | 1 | 0 | 1 | 1 | 1 | 1 | 1 | 3 | 1 |
| LD | animals with stripes | 2 | 1 | 2 | 2 | 0 | 1 | 2 | 0 | 1 | 2 | 0 | 1 | 0 | 2 |
| LD | long thin objects | 0 | 0 | 2 | 2 | 1 | 1 | 1 | 1 | 0 | 2 | 2 | 1 | 2 | 2 |
| LD | small objects | 1 | 2 | 0 | 1 | 2 | 1 | 0 | 1 | 1 | 1 | 1 | 0 | 1 | 1 |
| LD | things made of wood | 1 | 1 | 1 | 3 | 2 | 1 | 2 | 1 | 0 | 1 | 2 | 2 | 0 | 0 |
| LD | things that are blue | 2 | 1 | 1 | 1 | 1 | 1 | 1 | 1 | 0 | 0 | 1 | 0 | 1 | 1 |
| LD | things that are brown | 1 | 2 | 1 | 0 | 3 | 1 | 2 | 2 | 2 | 1 | 2 | 0 | 1 | 1 |
| LD | things that are green | 1 | 0 | 1 | 1 | 0 | 2 | 1 | 1 | 2 | 2 | 1 | 1 | 0 | 1 |
| LD | things that are orange | 2 | 1 | 2 | 2 | 1 | 2 | 0 | 1 | 1 | 1 | 0 | 2 | 2 | 1 |
| LD | things that are red | 1 | 1 | 1 | 1 | 0 | 1 | 1 | 1 | 2 | 0 | 2 | 1 | 2 | 1 |
| LD | things that are round | 1 | 1 | 1 | 1 | 1 | 1 | 0 | 1 | 3 | 1 | 2 | 0 | 1 | 1 |
| LD | things that are soft | 0 | 0 | 2 | 0 | 2 | 1 | 1 | 2 | 1 | 0 | 2 | 2 | 1 | 2 |
| LD | things that are very large | 1 | 1 | 1 | 2 | 1 | 2 | 1 | 1 | 1 | 1 | 1 | 2 | 3 | 0 |
| LD | things that are white | 3 | 2 | 1 | 1 | 1 | 0 | 1 | 0 | 1 | 1 | 0 | 3 | 0 | 2 |
| LD | things that are yellow | 2 | 3 | 0 | 0 | 0 | 1 | 2 | 2 | 1 | 3 | 1 | 1 | 2 | 0 |
| LD | things with doors | 0 | 1 | 1 | 0 | 1 | 1 | 2 | 2 | 1 | 1 | 0 | 1 | 2 | 3 |
| LD | things with handles | 0 | 1 | 1 | 1 | 2 | 1 | 1 | 1 | 1 | 1 | 1 | 1 | 0 | 0 |
