## Appendix 2 for "No evidence for a special role of language in feature-based categorization"

### Appendix 2: Experiment 3, fROI-Specific Results

Condition contrasts were designed to test the following null hypotheses.

Language network:

(1) $\frac{HD+LD}{2}=0$
(2) $LD=HD$	(main)
(3) $\frac{HD+LD2}{2}=Sentences$
(4) $\frac{HD+LD}{2}=Nonwords$

LD – low-dimensional categorization; HD – high-dimensional categorization.

Multiple demand network:

(1) $\frac{HD+LD}{2}=0$
(2) $LD=HD$	(main)
(3) $HardWM=EasyWM$
(4) $\frac{HD+LD}{2}=\frac{HardWM+EasyWM}{2}$
(5) $\frac{HD+LD}{2}=Sentences$
(6) $\frac{HD+LD}{2}=Nonwords$

HardWM – hard working memory task, EasyWM – easy working memory task.

Putative LD categorization regions (results reported in the main text):

(1) $\frac{HD+LD}{2}=0$
(2) $LD=HD$	(main)
(3) $HardWM=EasyWM$
(4) $Sentences=Nonwords$
(5) $\frac{HD+LD}{2}=\frac{HardWM+EasyWM}{2}$
(6) $\frac{HD+LD}{2}=Nonwords$

***Table 1****. Mixed-effect linear regression results for language fROIs. p-values were FDR-corrected for the number of fROIs. Significant p-values are highlighted in bold. S – sentence reading, N – nonword reading, LD – low-dimensional categorization, HD – high-dimensional categorization.*

| **ROI** | **Regression Term** | ***Beta*** | ***p-*value** |
| --- | --- | --- | --- |
| IFGorb | **Categorization>0** | **0.48** | **.005** |
|  | LD>HD | 0.04 | .932 |
|  | **S>Categorization** | **1.21** | **<.001** |
|  | N>Categorization | 0.02 | .905 |
| IFG | **Categorization>0** | **0.90** | **<.001** |
|  | LD>HD | 0.02 | .932 |
|  | **S>Categorization** | **1.41** | **<.001** |
|  | N>Categorization | -0.15 | .510 |
| MFG | **Categorization>0** | **0.63** | **.005** |
|  | LD>HD | 0.11 | .932 |
|  | **S>Categorization** | **2.41** | **<.001** |
|  | **N>Categorization** | **0.75** | **.031** |
| PostTemp | Categorization>0 | 0.27 | .094 |
|  | LD>HD | 0.10 | .932 |
|  | **S>Categorization** | **1.81** | **<.001** |
|  | N>Categorization | 0.26 | .114 |
| AntTemp | Categorization>0 | -0.02 | .760 |
|  | LD>HD | -0.08 | .932 |
|  | **S>Categorization** | **1.52** | **<.001** |
|  | N>Categorization | 0.22 | .114 |
| AngG | Categorization>0 | 0.24 | .165 |
|  | LD>HD | -0.30 | .575 |
|  | **S>Categorization** | **0.57** | **<.001** |
|  | N>Categorization | -0.34 | .086 |

***Table 2****. Mixed-effect linear regression results for multiple demand fROIs. P-values were FDR-corrected for the number of fROIs. Significant p-values are highlighted in bold. H – hard working memory task, E – easy working memory task, LD – low-dimensional categorization, HD – high-dimensional categorization.*

| **Hemisphere** | **fROI #** | **fROI name** | **Regression Term** | ***Beta*** | ***p*-value** |
| --- | --- | --- | --- | --- | --- |
| L | 1 | postParietal | **Categorization>0** | **1.44** | **<.001** |
|  |  |  | LD>HD | 0.28 | .902 |
|  |  |  | **Hard WM>Easy WM** | **1.37** | **<.001** |
|  |  |  | **WM>Categorization** | **2.69** | **<.001** |
|  |  |  | **Nonwords>Categorization** | **-0.90** | **<.001** |
|  |  |  | **Sentences>Categorization** | **-1.36** | **<.001** |
| L | 2 | midParietal | **Categorization>0** | **0.98** | **.004** |
|  |  |  | LD>HD | 0.30 | .902 |
|  |  |  | **Hard WM>Easy WM** | **1.40** | **<.001** |
|  |  |  | **WM>Categorization** | **2.32** | **<.001** |
|  |  |  | Nonwords>Categorization | -0.07 | .913 |
|  |  |  | **Sentences>Categorization** | **-0.65** | **.034** |
| L | 3 | antParietal | **Categorization>0** | **0.88** | **.003** |
|  |  |  | LD>HD | 0.47 | .902 |
|  |  |  | **Hard WM>Easy WM** | **1.14** | **<.001** |
|  |  |  | **WM>Categorization** | **2.10** | **<.001** |
|  |  |  | Nonwords>Categorization | -0.17 | .567 |
|  |  |  | **Sentences>Categorization** | **-0.71** | **.004** |
| L | 4 | supFrontal | **Categorization>0** | **0.77** | **.007** |
|  |  |  | LD>HD | 0.23 | .902 |
|  |  |  | **Hard WM>Easy WM** | **0.97** | **.002** |
|  |  |  | **WM>Categorization** | **2.16** | **<.001** |
|  |  |  | Nonwords>Categorization | -0.28 | .355 |
|  |  |  | Sentences>Categorization | -0.38 | .139 |
| L | 5 | precentral_A | **Categorization>0** | **2.23** | **<.001** |
|  |  |  | LD>HD | 0.37 | .902 |
|  |  |  | **Hard WM>Easy WM** | **1.34** | **<.001** |
|  |  |  | **WM>Categorization** | **0.98** | **<.001** |
|  |  |  | Nonwords>Categorization | -0.56 | .058 |
|  |  |  | **Sentences>Categorization** | **-0.97** | **<.001** |
| L | 6 | precentral_B | **Categorization>0** | **1.42** | **<.001** |
|  |  |  | LD>HD | 0.17 | .902 |
|  |  |  | **Hard WM>Easy WM** | **1.12** | **<.001** |
|  |  |  | **WM>Categorization** | **0.71** | **.001** |
|  |  |  | **Nonwords>Categorization** | **-0.66** | **.030** |
|  |  |  | **Sentences>Categorization** | **-0.77** | **.004** |
| L | 7 | midFrontal | **Categorization>0** | **1.29** | **<.001** |
|  |  |  | LD>HD | 0.18 | .902 |
|  |  |  | **Hard WM>Easy WM** | **0.94** | **<.001** |
|  |  |  | WM>Categorization | 0.25 | .178 |
|  |  |  | **Nonwords>Categorization** | **-0.86** | **.002** |
|  |  |  | **Sentences>Categorization** | **-1.25** | **<.001** |
| L | 8 | midFrontalOrb | **Categorization>0** | **1.15** | **.006** |
|  |  |  | LD>HD | 0.13 | .902 |
|  |  |  | **Hard WM>Easy WM** | **1.29** | **<.001** |
|  |  |  | **WM>Categorization** | **0.65** | **.003** |
|  |  |  | **Nonwords>Categorization** | **-0.62** | **.034** |
|  |  |  | **Sentences>Categorization** | **-1.08** | **<.001** |
| L | 9 | insula | **Categorization>0** | **0.81** | **<.001** |
|  |  |  | LD>HD | -0.01 | .953 |
|  |  |  | **Hard WM>Easy WM** | **0.72** | **<.001** |
|  |  |  | **WM>Categorization** | **0.49** | **<.001** |
|  |  |  | **Nonwords>Categorization** | **-0.38** | **.008** |
|  |  |  | **Sentences>Categorization** | **-0.49** | **<.001** |
| L | 10 | medialFrontal | **Categorization>0** | **0.95** | **<.001** |
|  |  |  | LD>HD | -0.01 | .953 |
|  |  |  | **Hard WM>Easy WM** | **0.79** | **<.001** |
|  |  |  | **WM>Categorization** | **0.59** | **<.001** |
|  |  |  | **Nonwords>Categorization** | **-0.40** | **.033** |
|  |  |  | **Sentences>Categorization** | **-0.56** | **.001** |
| R | 1 | postParietal | **Categorization>0** | **1.15** | **<.001** |
|  |  |  | LD>HD | 0.26 | .902 |
|  |  |  | **Hard WM>Easy WM** | **1.68** | **<.001** |
|  |  |  | **WM>Categorization** | **3.18** | **<.001** |
|  |  |  | **Nonwords>Categorization** | **-0.84** | **.009** |
|  |  |  | **Sentences>Categorization** | **-1.14** | **<.001** |
| R | 2 | midParietal | **Categorization>0** | **0.74** | **.004** |
|  |  |  | LD>HD | 0.31 | .902 |
|  |  |  | **Hard WM>Easy WM** | **1.72** | **<.001** |
|  |  |  | **WM>Categorization** | **2.09** | **<.001** |
|  |  |  | Nonwords>Categorization | 0.05 | .913 |
|  |  |  | Sentences>Categorization | -0.43 | .148 |
| R | 3 | antParietal | **Categorization>0** | **0.33** | **.040** |
|  |  |  | LD>HD | 0.35 | .902 |
|  |  |  | **Hard WM>Easy WM** | **1.23** | **<.001** |
|  |  |  | **WM>Categorization** | **1.98** | **<.001** |
|  |  |  | Nonwords>Categorization | 0.00 | .989 |
|  |  |  | Sentences>Categorization | -0.27 | .273 |
| R | 4 | supFrontal | **Categorization>0** | **0.70** | **.013** |
|  |  |  | LD>HD | 0.14 | .902 |
|  |  |  | **Hard WM>Easy WM** | **1.55** | **<.001** |
|  |  |  | **WM>Categorization** | **2.77** | **<.001** |
|  |  |  | Nonwords>Categorization | -0.06 | .913 |
|  |  |  | Sentences>Categorization | -0.20 | .526 |
| R | 5 | precentral_A | **Categorization>0** | **1.48** | **<.001** |
|  |  |  | LD>HD | 0.19 | .902 |
|  |  |  | **Hard WM>Easy WM** | **1.40** | **<.001** |
|  |  |  | **WM>Categorization** | **1.22** | **<.001** |
|  |  |  | Nonwords>Categorization | -0.35 | .353 |
|  |  |  | **Sentences>Categorization** | **-0.68** | **.032** |
| R | 6 | precentral_B | **Categorization>0** | **1.73** | **<.001** |
|  |  |  | LD>HD | 0.29 | .902 |
|  |  |  | **Hard WM>Easy WM** | **1.65** | **<.001** |
|  |  |  | **WM>Categorization** | **1.21** | **<.001** |
|  |  |  | Nonwords>Categorization | -0.72 | .067 |
|  |  |  | **Sentences>Categorization** | **-1.08** | **.004** |
| R | 7 | midFrontal | **Categorization>0** | **0.91** | **.009** |
|  |  |  | LD>HD | 0.17 | .902 |
|  |  |  | **Hard WM>Easy WM** | **1.71** | **<.001** |
|  |  |  | **WM>Categorization** | **1.08** | **<.001** |
|  |  |  | Nonwords>Categorization | -0.26 | .485 |
|  |  |  | **Sentences>Categorization** | **-0.64** | **.036** |
| R | 8 | midFrontalOrb | **Categorization>0** | **0.89** | **.003** |
|  |  |  | LD>HD | 0.05 | .953 |
|  |  |  | **Hard WM>Easy WM** | **1.89** | **<.001** |
|  |  |  | **WM>Categorization** | **0.96** | **<.001** |
|  |  |  | Nonwords>Categorization | -0.39 | .306 |
|  |  |  | **Sentences>Categorization** | **-0.86** | **.007** |
| R | 9 | insula | **Categorization>0** | **0.72** | **.001** |
|  |  |  | LD>HD | -0.04 | .902 |
|  |  |  | **Hard WM>Easy WM** | **0.85** | **<.001** |
|  |  |  | **WM>Categorization** | **0.42** | **<.001** |
|  |  |  | **Nonwords>Categorization** | **-0.34** | **.013** |
|  |  |  | **Sentences>Categorization** | **-0.46** | **<.001** |
| R | 10 | medialFrontal | **Categorization>0** | **0.84** | **<.001** |
|  |  |  | LD>HD | 0.06 | .902 |
|  |  |  | **Hard WM>Easy WM** | **1.23** | **<.001** |
|  |  |  | **WM>Categorization** | **0.63** | **<.001** |
|  |  |  | Nonwords>Categorization | -0.35 | .073 |
|  |  |  | **Sentences>Categorization** | **-0.60** | **.002** |
